# Single cell analysis reveals an antiviral network that controls Zika virus infection in human dendritic cells

**DOI:** 10.1101/2024.01.19.576293

**Authors:** Kathryn M. Moore, Adam-Nicolas Pelletier, Stacey Lapp, Amanda Metz, Gregory K. Tharp, Michelle Lee, Swati Sharma Bhasin, Manoj Bhasin, Rafick-Pierre Sékaly, Steven E. Bosinger, Mehul S. Suthar

## Abstract

Zika virus (ZIKV) is a mosquito-borne flavivirus that caused an epidemic in the Americas in 2016 and is linked to severe neonatal birth defects, including microcephaly and spontaneous abortion. To better understand the host response to ZIKV infection, we adapted the 10x Genomics Chromium single cell RNA sequencing (scRNA-seq) assay to simultaneously capture viral RNA and host mRNA. Using this assay, we profiled the antiviral landscape in a population of human moDCs infected with ZIKV at the single cell level. The bystander cells, which lacked detectable viral RNA, expressed an antiviral state that was enriched for genes coinciding predominantly with a type I interferon (IFN) response. Within the infected cells, viral RNA negatively correlated with type I IFN dependent and independent genes (antiviral module). We modeled the ZIKV specific antiviral state at the protein level leveraging experimentally derived protein-interaction data. We identified a highly interconnected network between the antiviral module and other host proteins. In this work, we propose a new paradigm for evaluating the antiviral response to a specific virus, combining an unbiased list of genes that highly correlate with viral RNA on a per cell basis with experimental protein interaction data. Our ZIKV-inclusive scRNA-seq assay will serve as a useful tool to gaining greater insight into the host response to ZIKV and can be applied more broadly to the flavivirus field.

## Introduction

Zika virus (ZIKV) is a mosquito-borne flavivirus that caused an epidemic in the Americas in 2016. Other routes of transmission include sexual intercourse, vertical transmission from mother to fetus in utero, and blood transfusion. ZIKV was first isolated in Uganda in 1947 (1) and remained dormant in Africa and Asia for decades, with sporadic outbreaks characterized by a mild self-limiting disease in humans (2). The emergence of ZIKV in the Americas corresponded with enhanced disease severity, characterized by severe neonatal birth defects, including microcephaly and spontaneous abortion, and Guillain-Barre syndrome in adults (3).

Dendritic cells (DCs) are critical for protective immunity against several mosquito-borne flaviviruses, including West Nile virus (WNV) (4–6), Dengue virus (DENV) (7–9), and Yellow Fever virus (YFV) (10, 11). Despite their antiviral activity, recent studies in humans and non-human primates have found that DCs are early target cells of ZIKV infection (12–16). Human monocyte derived dendritic cells (moDCs) have been used as a model to study the host response to ZIKV infection (17–22). We previously found that contemporary and historic ZIKV strains replicate in DCs (19). Yet ZIKV infected DCs failed to stimulate secretion of proinflammatory cytokines or induce T cell proliferation (18, 19). These trends have been observed in clinical samples in which DCs isolated from ZIKV-infected patients revealed downregulation in transcriptional pathways related to phagocytosis and antigen presentation suggesting ZIKV mediated immune evasion (23, 24).

The type I interferon (IFN) response is critical in host defense against flavivirus infection. ZIKV is recognized by the RIG-I like receptors (RLR), RIG-I and MDA5, which activate the transcription factors (IRF-3/7 and NF-kB), leading to production of type I IFN (IFN-α/IFN-β). Type I IFN signals in an autocrine and paracrine manner through the IFN-α/β receptor (IFNAR). This leads to a signaling cascade via STAT molecules that results in expression of hundreds of IFN stimulated genes (ISGs), some of which have antiviral activity. ZIKV inhibits type I IFN downstream signaling at multiple points along the pathway such as inhibition of STAT signaling via blocking JAK1 and TYK2 activation and STAT2 degradation, allowing it to evade host restriction (20, 25–32). However, in past studies with primary human cells, we have still observed high levels of ISG expression in ZIKV infected cell cultures (19, 33, 34).

There is growing evidence that virus infected cells and the uninfected bystander cells have phenotypic differences such as differences in expression of activation markers, cytokines and genes related to innate immune response and metabolic pathways (19, 22, 35–40). Virus-inclusive single cell RNA sequencing (scRNA-seq) is a single cell approach that allows for detection of viral RNA and host transcripts within the same cell. This technique allows for distinguishing the transcriptional profiles of infected and bystander cells, as well as detect other subpopulations and cell-to-cell heterogeneity that are masked in bulk assays (37, 41–45). There have been a few investigations using scRNA-seq to understand ZIKV infection in other cell types such as neural stem and progenitor cells (46–48), Huh7 cells (49), and testicular tissue (50). Yet, the impact that ZIKV has on the antiviral response at the single cell level in DCs has yet to be explored (47, 49).

Since most scRNA-seq assays use poly-dT primers to capture polyadenylated mRNA and flaviviruses do not have polyadenylated RNA, virus-specific primers have been added to the assay to detect viral RNA from other flaviviruses alongside host mRNA (43, 49, 51–53). However, these prior assays were based on scRNA-seq platforms reliant on index sorting the cells, which limits the number of cells that can be easily profiled. In this study, we developed a ZIKV-inclusive scRNA-seq protocol using the 10x Genomics Chromium assay, a higher throughput assay than prior flavivirus-inclusive assays (54). We then applied this assay to define the transcriptional signatures of ZIKV infected and bystander cells in a population of human moDCs with a low level of infected cells to model early viral infection. Bystander cells expressed an antiviral state that was enriched for genes coinciding with a type I IFN response. Within the infected cells, viral RNA negatively correlated with type I IFN dependent and independent genes. Using experimentally derived protein-interaction data, we found that these genes that highly correlate with viral RNA on a per cell basis form an interconnected network on the protein level. Here we propose that the antiviral response to ZIKV is a coordinated network of host genes that collectively restrict viral RNA, rather than a collection of antiviral effectors acting in parallel.

## Results

### ZIKV-specific primer enhances detection of viral RNA using single cell RNA sequencing

In this study, we used the 10x Genomics Chromium Next GEM Single Cell 5’ assay, which uses a microfluidic chip to generate a water-in-oil emulsion (termed GEMs) with each water droplet containing a cell, the reagents for reverse transcription, and a unique DNA barcode (**Figure 1**). A poly-dT primer is used to initiate reverse transcription of polyadenylated mRNA for sequencing. Since the ZIKV genome does not have a polyadenylated tail, we incorporated a ZIKV-specific primer into the assay (**Figure 1A**). The ZIKV-specific primer used in this study was previously used as a laboratory diagnostic assay and was modified to contain a non-poly-dT tag for amplification along with host cDNA after reverse transcription (55) (**Figure 1B**). We determined the optimal concentration of the ZIKV-specific primer in VeroE6 cells. The lowest concentration tested (0.1 µM) maximized detection of viral RNA without compromising detection of host mRNAs and proceeded with this concentration throughout the rest of the study (**Supplemental Figure 1**).

**Figure 1.**
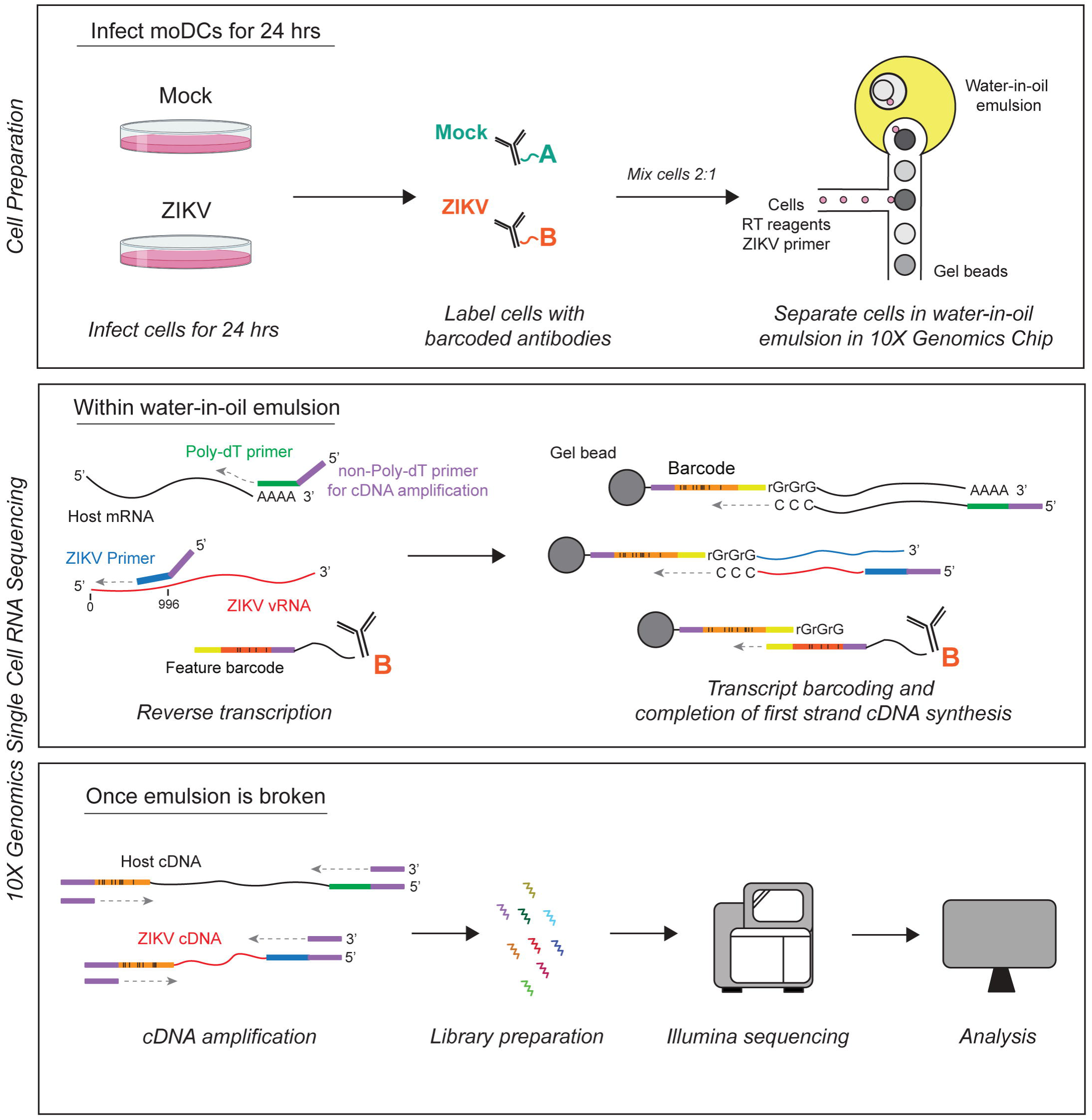
Schematic for ZIKV-inclusive single cell RNA sequencing using the 10x Genomics assay. Overview of the cell preparation and 10x Genomics Chromium scRNA-seq assay. Mock and ZIKV treated cells are labeled with Feature barcoded antibodies, mixed 2:1, and then loaded in the 10x microfluidic chip with the ZIKV-specific primer. The reverse transcription reaction of host mRNA, ZIKV viral RNA, and Feature barcode takes place in individual water-in-oil emulsion droplets (GEMs). A unique barcode is added onto the nascent cDNA, allowing each transcript to be computationally traced back to its original cell. Created with BioRender.com.

We next applied the ZIKV-inclusive assay using moDCs, which are target cells of infection and have been extensively used to model the host response to infection (12–21). Monocytes were isolated from human blood and differentiated into moDCs using GM-CSF and IL-4 (19, 20). moDCs were treated with ZIKV (PRVABC59) at an MOI 2.5 and prepared for either flow cytometry or scRNA-seq (**Figure 2**). At this MOI, ZIKV was found to infect 3.22% of cells at 24 hpi as demonstrated by ZIVK E protein staining by flow cytometry (**Figure 2A, Supplemental Figure 1**). Presence of ambient viral RNA released into the media prior to separation into single cell droplets has been observed by others during droplet-based scRNA-seq assays (56). Extracellular viral RNA within the water droplet of an uninfected bystander cell may lead to viral RNA reads being sequenced and may reduce confidence in identifying infected cells. We used Feature Barcoding to computationally separate cell associated ZIKV viral RNA from ambient ZIKV viral RNA found within a cell suspension. Mock and ZIKV treated cells were surface labeled with an antibody containing a feature barcode (termed either Hash A or Hash B in **Figure 2B**) with separate sequences from the cell specific barcodes and then mixed prior to capture 2:1. These barcodes were reverse transcribed and barcoded with the cell-specific barcode along with host mRNA and viral RNA. After sequencing, the cells were computationally identified as either mock (Hash A) or ZIKV treated (Hash B). Ambient RNA was measured as any detected viral RNA in the mock cells. We found that in the ZIKV-inclusive assay, 89% of mock cells had 0 normalized viral RNA reads, leaving 11% of the cells containing some detectable amount of ambient viral RNA (**Figure 2C**). Only 1.36% of mock cells had normalized viral RNA reads of >1 and the majority of mock cells contaminated with ambient viral RNA had low levels, with normalized expression between 0-1. Based on these results, we considered a ZIKV treated cell with expression greater than 1 to be likely infected. The threshold for infection was set at 1; Hash B labeled cells treated with ZIKV for 24 hours with normalized viral RNA reads ≥1 labeled as “infected” cells and those with normalized viral RNA reads <1 labeled “bystander.” We could not confidently distinguish infected cells at a viral RNA range of 0-1, thus these cells were excluded from downstream analysis. Hash A labeled cells with 0 normalized viral RNA reads were labeled “mock” and all other cells were excluded. A UMAP of the final data set was created and separated based on those annotations, where each point represents a single cell (**Figure 2C**). In the sample without the ZIKV-specific primer, 1.69% of mock cells had normalized viral RNA reads between 0 and 1, compared to 9.64% in the ZIKV-inclusive group. Conversely, more cells had undetectable viral RNA in the sample without the ZIKV-specific primer compared to samples with the ZIKV-specific primer.

**Figure 2.**
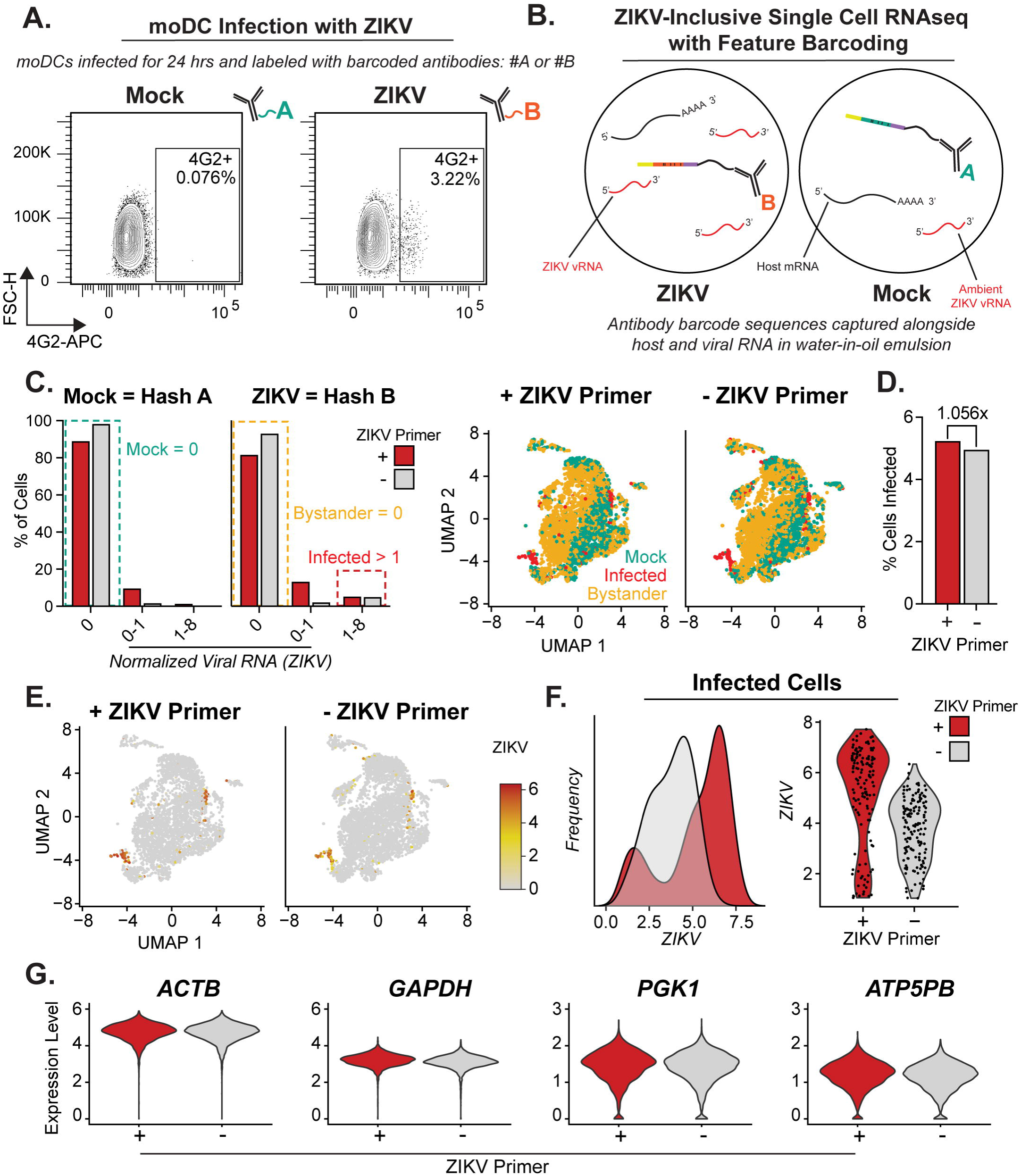
ZIKV-inclusive assay was used in moDCs to distinguish between infected and bystander cells. Primary human monocyte derived dendritic cells (moDCs) were infected with ZIKV and the ZIKV-inclusive scRNA-seq assay was performed at 24 hpi. (A) Flow cytometry plot of mock and ZIKV-infected cells stained for ZIKV E protein expression using the pan-flavivirus antibody, 4G2. (B) Mock and ZIKV-infected cells were labeled with feature barcoded antibodies, Hash A and Hash B, respectively. They were then mixed 2:1. These barcodes were captured during the reverse transcription reactions alongside host mRNA allowing for mock and ZIKV cells to be distinguished within the same sample computationally. (C) Distribution of normalized viral RNA reads across Hash A (mock) and Hash B (ZIKV) represented as percent of cells in samples with and without ZIKV-specific primer. Cells were labeled as mock (Hash A, ZIKV = 0), bystander (Hash B, ZIKV = 0), and infected (Hash B, ZIKV > 1) and color coded as green, yellow, and red, respectively in the dashed lines on the bar plot (left) and as the dots on the UMAP (right). (D) Percent of cells in ZIKV population (Hash B) that were determined to be infected in samples with and without ZIKV-specific primer. (E) UMAPs of samples with and without ZIKV-specific primer featuring viral RNA. (F) Distribution of normalized viral RNA reads in infected cells across samples with and without ZIKV-specific primer represented as a histogram (left) and violin plot (right). (G) Expression of housekeeping genes across samples with and without ZIKV-specific primer. All graphs comparing samples with and without ZIKV-specific primer are colored red (+ ZIKV-specific primer) and grey (-ZIKV-specific primer).

Detection of ZIKV-infected cells was compared in samples with and without ZIKV-specific primer. There was a 5.6% increase in the percent of cells identified as infected with the addition of the ZIKV-specific primer (5.23% infected) compared to the sample without a ZIKV-specific primer (4.96% infected) (**Figure 2D**). The range of viral RNA in infected cells was expanded in the sample containing ZIKV-specific primer (**Figure 2E-F**). In the ZIKV-inclusive assay, we observed two populations of infected cells, those with low expression and high expression, that are not observed in the samples without the ZIKV-specific primer (**Figure 2F**). Similar to our observations in VeroE6 cells, housekeeping gene expression (*ACTB*, *GAPDH*, *PGK1*, *ATP5PB*) was similar between the two groups, indicating no detrimental cost of adding the additional primer into the reverse transcription reaction (**Figure 2G**). These findings suggest that including the ZIKV-specific primer enhanced detection of low-level viral RNA and in the sample without the ZIKV-specific primer, any potentially lowly infected cells are left undetected and labeled as bystander cells.

### Transcriptional signatures in ZIKV-infected and bystander cells are distinct

ZIKV-infected cells and bystander cells were compared to mock and analyzed for enrichment in Hallmark gene sets by Gene Set Enrichment Analysis (**Figure 3**). Enrichment scores in infected and bystander (relative to mock) were compared and found to be divergent in many of these biological pathways (**Figure 3A**). Pathways related to viral infection such as IFN alpha/gamma responses, TNFA signaling via NF-κB, and inflammatory response were enriched in bystander cells when compared to mock (p-value < 0.05) and the infected cells were not. We also observed that IL2-STAT5 and IL6-JAK-STAT3 signaling pathways were negatively enriched in infected cells, meaning that expression of genes in those pathways was higher in uninfected (mock) cells, whereas the bystander cells were positively enriched.

**Figure 3.**
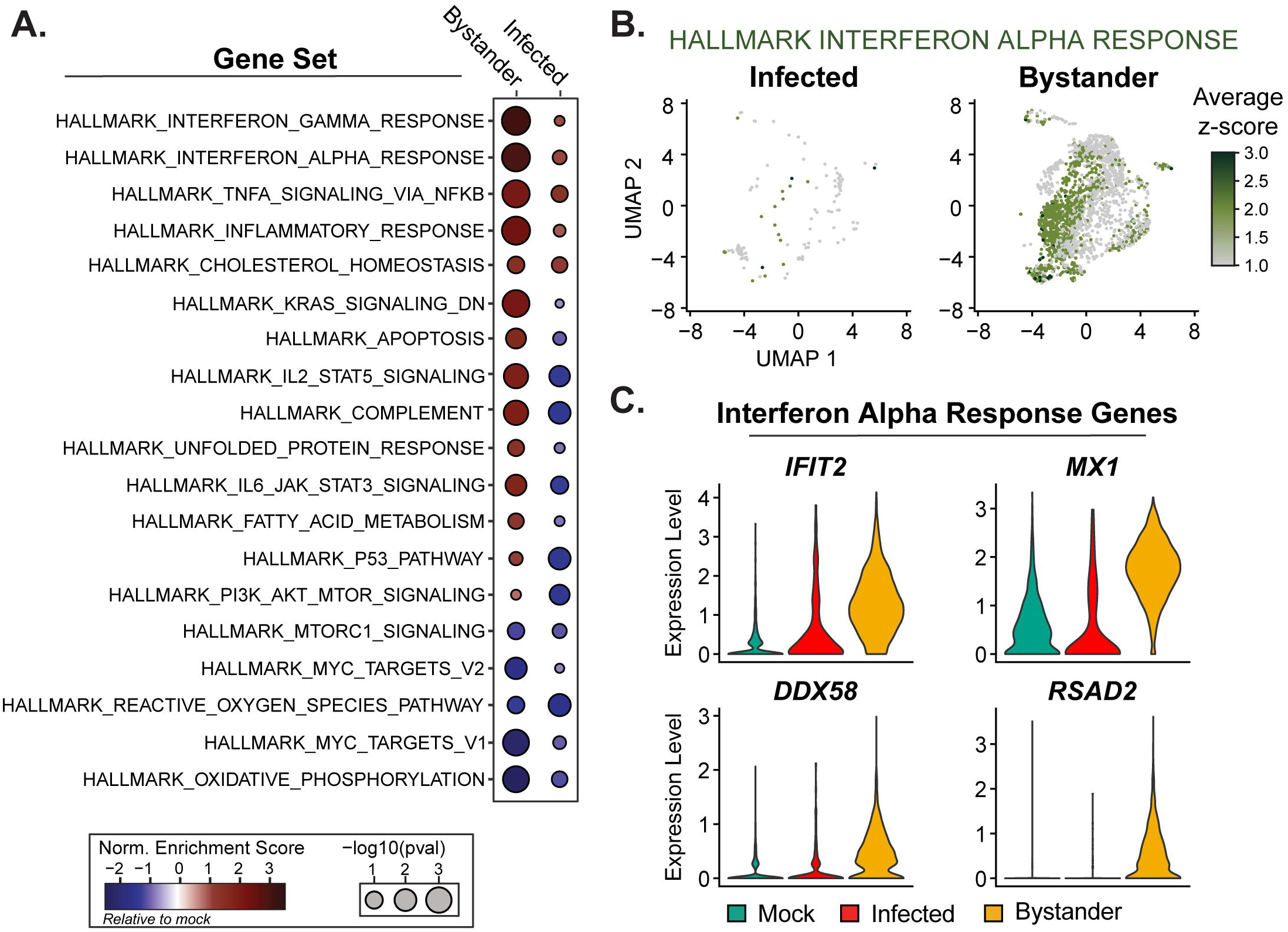
ZIKV-infected and bystander moDCs have diverging enrichment in key antiviral pathways. (A) Gene set enrichment analysis using Hallmark gene sets. Size of circles represents −log10 p-value and color bar represents normalized enrichment score of infected or bystander cells relative to mock cells. (B) UMAPs featuring the average z-score for Hallmark IFN Alpha Response pathway. (C) Expression of specific genes in Hallmark IFN Alpha Response pathway.

UMAPs featuring enrichment in the Hallmark IFN Alpha Response pathway illustrates heterogeneity across individual moDCs in both infected and bystander cells (**Figure 3B**). Further analysis showed that the expression of specific genes from the pathway such as *IFIT2, MX1, DDX58, RSAD2*, were similar between mock and infected, yet it was the bystander cells that were upregulating expression (**Figure 3C**).

### IFN-β expression is rare and restricted to ZIKV-infected cells

It has previously been reported that there is a low number of IFN expressing cells in a population of cells infected with other viruses (43, 57–59). We also observed a small number of cells expressing type I IFN gene expression after exposure to ZIKV (**Figure 4**, **Figure 4A**). *IFNA1* expression was observed in mock, ZIKV-infected, and bystander cells, but ZIKV-infected cells had a higher rate (15 per 1,000 cells) compared to the mock and bystander (<1 per 1,000 cells) (**Figure 4B**). *IFNB1* expression was exclusively seen in infected cells and was expressed at a rate of 51 per 1,000 infected cells (5%) and 3 per 1,000 cells in the total population (0.3%). Although we did not observe a trend in *IFNB1* expression with viral RNA, all cells that expressed *IFNB1* had higher levels of viral RNA (>3.5, **Figure 4C**). We observed a clear bimodal distribution of viral RNA across the infected cells and indeed, the *IFNB1* expressing cells fell within population of high viral RNA containing infected cells (**Figure 4C-D**).

**Figure 4.**
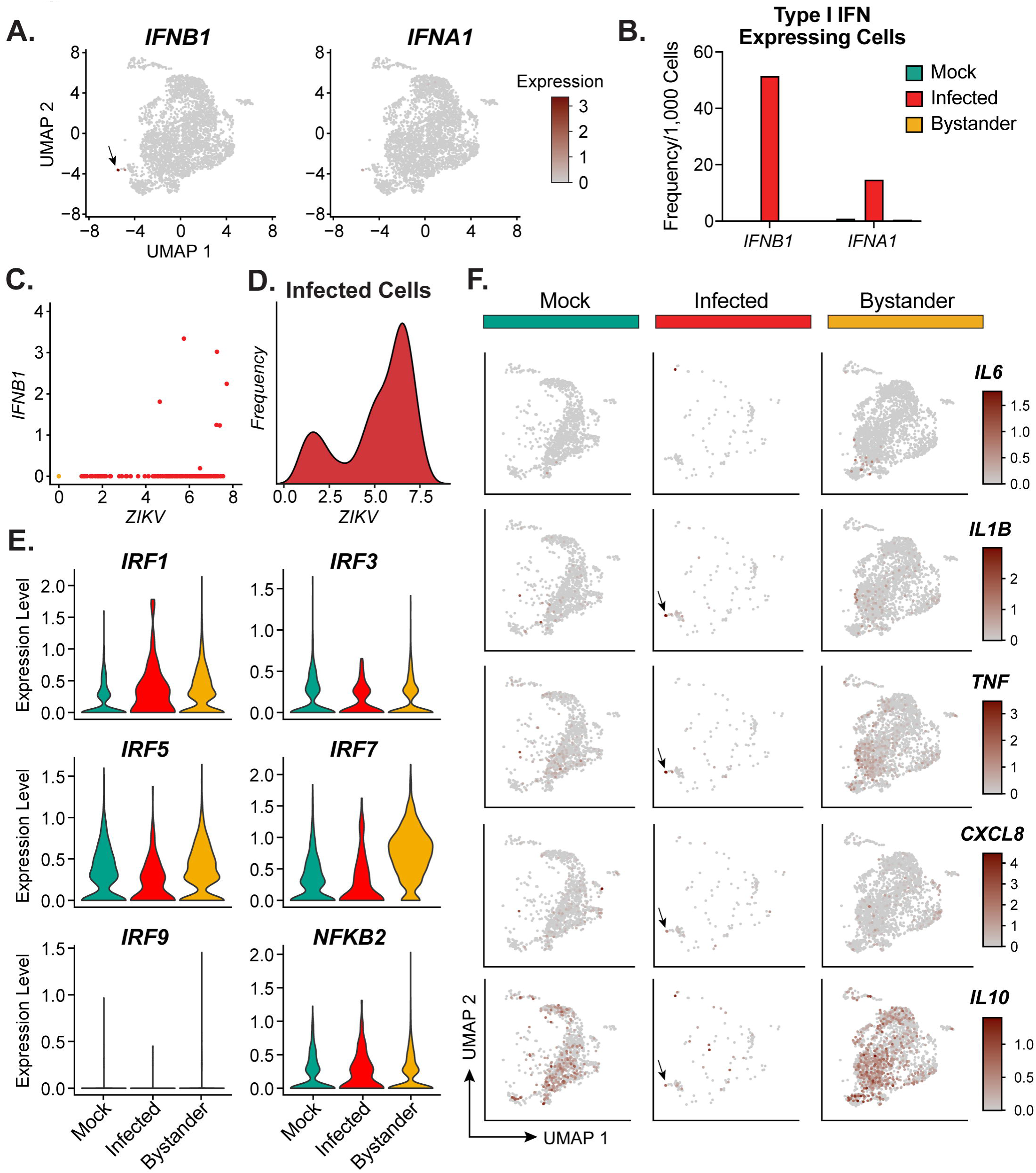
Induction of Type I IFN in moDCs exposed to ZIKV. (A) UMAP featuring IFN expression in sample containing mock, infected, and bystander cells. (B) Frequency of cells expressing type I IFN within each group. (C) Scatter plot between *ZIKV* viral RNA and *IFNB1* expression for each cell, labeled with the same colors as in B. (D) Bimodal distribution of viral RNA among infected cells. (E) Expression of IRFs (*IRF1*, *IRF3*, *IRF5*, *IRF7*, *IRF9*) and *NFKB2* transcription factors across mock, infected, and bystander cells. (F) UMAPs split across groups featuring expression of inflammatory (*IL6*, *IL1B*, *TNF*, *CXCL8*) and anti-inflammatory (*IL10*) cytokines.

Type I IFN is induced upon viral RNA sensing by RLRs via transcription factors such as IFN regulatory factors (IRFs) and NF-κB. To better understand the potential basis for type I IFN induction in the context of ZIKV infection, we evaluated the expression of IRF and NF-κB genes across all groups (**Figure 4E**). Notably, the trends in IRF expression between groups were not consistent across all IRFs. While ZIKV did induce type I IFN expression, *IRF1* was the only IRF that was increased in infected cells. There was also a marginal increase in *NFKB2* expression in infected cells as well. This was in contrast to *IRF7*, the master regulator of type I IFN expression, which showed similar if not slightly reduced expression among infected cells compared to mock. However, there was a clear increase in *IRF7* expression among bystander cells, yet minimal type I IFN induction. To understand if the restricted induction of type I IFN was unique, we evaluated the expression of other cytokines (**Figure 4F**). Cells in the same UMAP region in the infected cells where *IFNB1* expression occurred (**Figure 4F, black arrows**) also expressed *IL1B*, *TNF*, *CXCL10*, and *IL10* suggesting that these *IFNB1* expressing cells also expressed other cytokines and held a similar transcriptional phenotype. However, expression of other inflammatory cytokines (*IL6*, *IL1B*, *TNF*, *CXCL8*) and the anti-inflammatory cytokine (*IL10*) were less restricted than *IFNB1*. There was an increase in number of cells expressing these other cytokines in the bystander cells, as well as baseline expression in some mock cells.

### Response to Type I IFN in moDCs is heterogeneous and occurs predominantly in bystander cells

To assess the transcriptional signature of IFN-β in moDCs, we included a sample treated with 100 IU/mL soluble human IFN-β for 24 hrs (**Figure 5**, **Figure 5A**). Differentially expressed genes (DEG) relative to mock were compared across infected, bystander, and IFN-β groups (**Figure 5B**). There was only 1 common DEG upregulated and 2 DEGs downregulated relative to mock among IFN-β treatment, ZIKV-infected, and ZIKV bystander cells (**Figure 4E**). There were more common DEGs between the IFN-β treated cells and bystander (100 and 8 genes up and down, respectively) than IFN-β treated and infected (4 and 16 genes up and down, respectively). There were only 6 total unique DEGs in bystander cells and 9 shared between infected and bystander. All other bystander DEGs (90%, 111/123) were common with IFN-β treatment, suggesting that transcriptional response in the bystanders is primarily a result of signaling from IFN-β-expressing infected cells. 19% of the DEGs in the IFN-β treatment group were induced in the bystander group. The upregulated genes in common between IFN-β treatment and bystander cells included *ISG15*, *IFIT1*, *IFI44*, *IFI6*, *IFITM3*, *OAS1*, *MX1*, *CXCL10*, and *RSAD2*. A total of 20 DEGs from ZIKV infection were shared with DEGs from IFN-β treatment, indicating some type I IFN response from ZIKV infection, but less prevalent than in bystander cells. ZIKV appeared to have a silencing effect on infected moDCs with 91% (111/122) total DEGs down-regulated relative to mock, in contrast to the bystander cells, whose DEGs were primarily up relative to mock (83%, 102/123).

**Figure 5.**
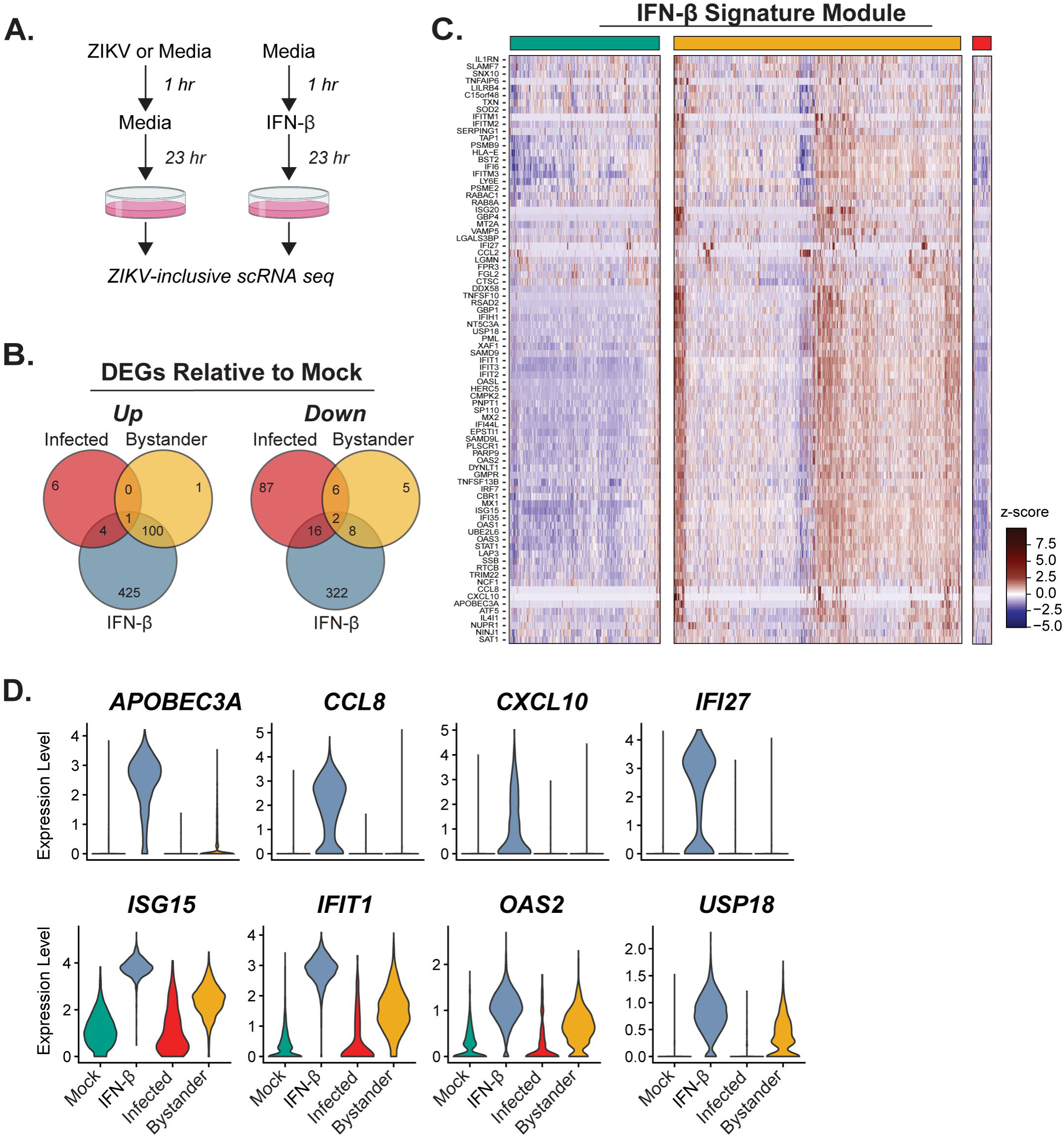
Type I IFN response in moDCs exposed to ZIKV. (A) Schematic for experiment comparing impact of IFN-β treatment after ZIKV infection moDCs. (B) Venn diagram of DEGs relative to mock in IFN-β treatment, infected, and bystander cells. DEGs were filtered based on adj p-value < 0.05 in each group relative to mock. (C) Expression of top DEGs within the IFN-β treatment (termed IFN-β module) in mock, bystander, and ZIKV-infected cells. IFN-β treatment DEGs were filtered based on adj p-value < 0.05 and average log2 fold change > 1 in IFN-β treatment relative to mock. Rows are genes and columns are individual cells organized by group. Data are represented as row z-scores. (D) Expression of representative genes from the IFN-β module across all groups. (E) Schematic for IFN-β treatment post-ZIKV infection. (F) Percent of infected cells in each sample. (G) Detected viral RNA across samples with and without IFN-β treatment post-infection. Colors of each group are consistent across the entire Figure: mock = green, IFN-β treatment = blue, infected = red, bystander = yellow. DEGs = differentially expressed genes.

The top DEGs that are expressed in the IFN-β group were termed the IFN-β module (**Figure 5C**). Bystander cells contained subpopulations of cells that scored high, medium, and low within the IFN-β module compared to mock or infected cells. Indeed, we observed heterogeneity in expression for the IFN-β module across all groups, but most prominently in bystander cells, which had the widest range of row z-scores (ranging from +10 to −5.03). Even within these subpopulations of bystander cells, on a per cell basis, there is substantial heterogeneity whereby not all genes are expressed within the IFN-β module. There were several genes in which expression was consistently low across the majority of cells with a few small clusters of cells that were very high expressing, including *APOBEC3A*, *CCL8*, *CXCL10*, and *IFI27* (**Figure 5D**). This contrasted with most other genes such as *ISG15*, *IFIT1*, *OAS2*, and *USP18,* which represent the most common expression pattern observed in the heatmap and that had more gradual distribution of expression across the cells. IFN-β treated cells also did not express the same genes or to the same degree, demonstrating that not all cells respond to IFN-β in a similar manner (**Supplemental Figure 2**).

### Antiviral transcriptional module associates with control of virus replication in moDCs

We screened all detectable host transcripts for correlation with viral RNA in infected cells (**Figure 6A**). There were 151 genes with strong negative correlation with viral RNA (Pearson’s correlation coefficients (ρ) < −0.3 and adj p-value < 0.05) which we defined as the antiviral module. We next compared the antiviral module (151 genes) to the IFN-β module (82 genes). There was substantial overlap between the antiviral and IFN-β modules, with 40 common genes between the antiviral module and the IFN-β module. This amounted to 49% (40/82) of IFN-β module genes strongly negatively correlated with viral RNA (Figure 6B). Heatmaps depicting the correlation coefficients of viral RNA and module genes ordered based on strength of negative correlation with viral RNA can be observed in **Figure 6C**. Genes from the antiviral module that highly negatively correlated with viral RNA were more likely to positively correlate with each other. The same trend was observed for the IFN-β module. Next, the average expression for the antiviral and IFN-β modules were calculated for each cell and plotted versus viral RNA in infected cells (**Figure 6D**). The IFN-β module score negatively correlated with viral RNA (ρ = −0.63) but lower compared to the antiviral module score (ρ = −0.82).

**Figure 6.**
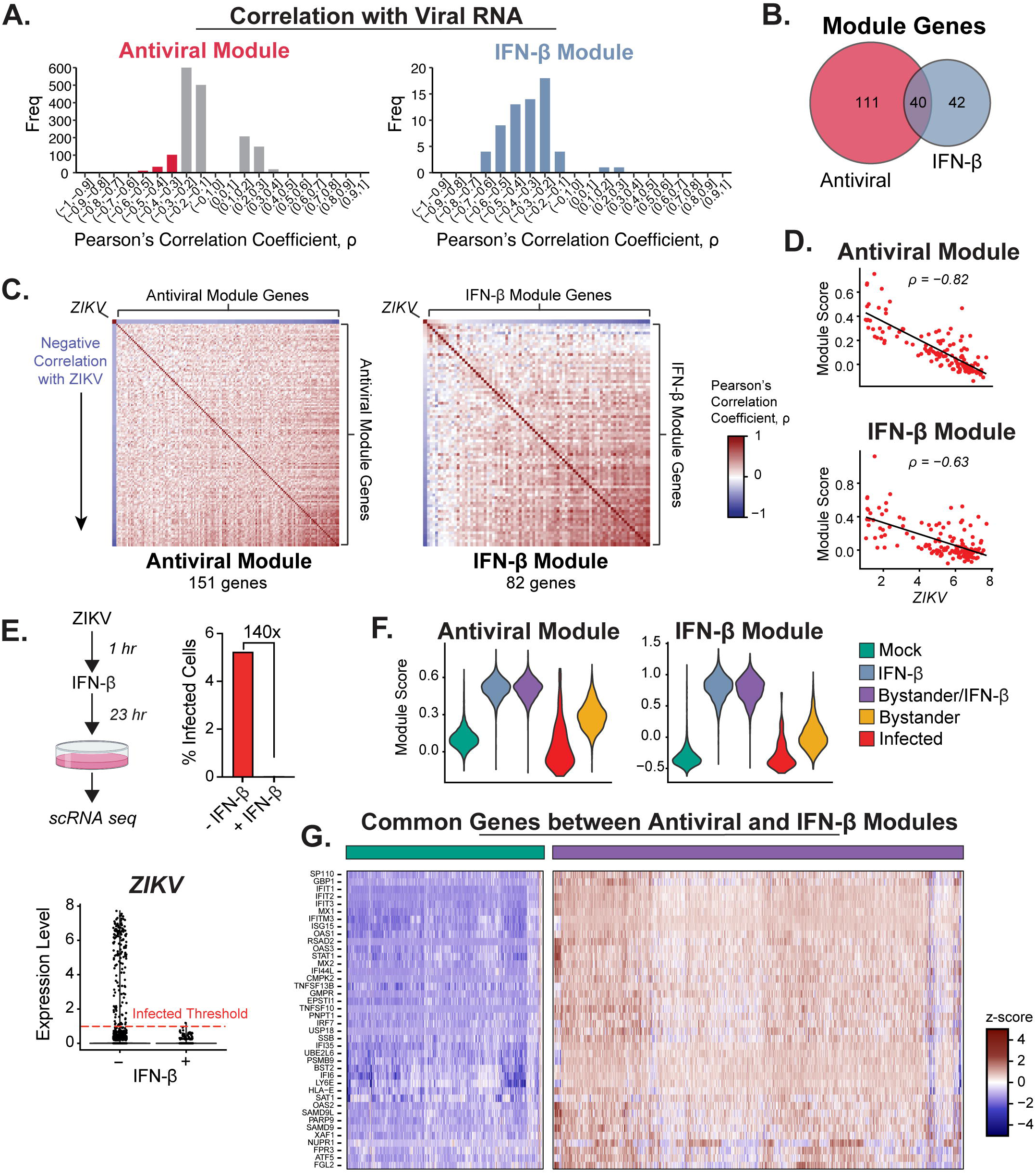
Analysis of ZIKV-specific antiviral module. (A-B) Frequency distribution of Pearson’s correlation coefficients (ρ) between viral RNA and host genes: (left) All significantly correlating host genes (filtered by p < 0.05) are displayed, with red bar indicating the frequency strong negative correlating genes that comprise the antiviral module (ρ < −0.3); (right) IFN-β module only. (B) Venn diagram showing overlap in genes between antiviral and IFN-β module. (C) Pearson’s correlation coefficients between ZIKV viral RNA and modules displayed as a heatmap, where genes are ordered based on strength of negative correlation with viral RNA. (D) Correlation between viral RNA and average expression of genes within each module (termed module score). (E) Schematic for ZIKV infection of moDCs followed by soluble human IFN-β treatment (left); percent of infected cells in each sample (right); viral RNA expression across groups (bottom). (F) Module score expression across all groups. (G) Heatmap of the 40 common genes between antiviral and IFN-β module expressed in mock (left, green) and bystander cells that received the IFN-β post-infection treatment (right, purple). Data are represented as row z-scores.

Given the substantial overlap between the antiviral and IFN-β modules, we treated cells with IFN-β 1 hour post-infection to functionally validate our antiviral module. As previously observed, IFN-β treatment reduced viral RNA and the percentage of infected cells (**Figure 6E**) (19, 29). We detected only a single cell that had ZIKV viral RNA above the threshold of a limit of detection (>1), which resulted in a 140x reduction in infected cells. We next wanted to investigate if IFN-β treatment post-infection resulted in enrichment in the antiviral and IFN-β modules. Indeed, we found that cells treated with IFN-β were almost identical to IFN-β treatment alone for both modules (**Figure 6F**). Infected and mock cells had similar module scores and were the lowest groups on average, whereas the uninfected bystander cells had elevated scores. Expression of the 40 overlapping genes that strongly correlate with viral RNA (antiviral module) were induced in cells with IFN-β treatment post-infection (**Figure 6G**). Notably, *IRF7* was one of the genes in both modules that correlated negatively with viral RNA. These results demonstrate that expression of genes within the IFN-β module, including a subset of genes, contributes to restricting ZIKV infection.

### Antiviral module proteins form an interaction network based on experimental data

We next sought to model the antiviral module at the protein level using existing protein-protein interaction datasets. Using open-source molecular interaction database IntAct, we extracted human protein-protein interaction data derived from experiments such as pull down, co-immunoprecipitation, and two-hybrid assays, indicating direct interactions or physical associations in complexes containing two or more proteins. Three of the 151 antiviral module genes encoded long non-coding RNAs and were excluded from the analysis. We found that the antiviral module proteins formed an extensive and complex protein-interaction network (**Figure 7**). Considering that some ISGs require interactions with other host proteins to become activated or carry out their function, we included all known interacting partners for each antiviral module protein (first neighbor interactors). There were known interactions between the antiviral module proteins as well as indirect interactions through a common first neighbor (one degree away). The antiviral module proteins with greatest number of edges and thus interacted with the greatest number of different proteins were PLEC (Plectin), SYK (Spleen Associated Tyrosine Kinase), EIF2AK2 (Protein Kinase R), STAT1 (Signal Transducer and Activator of Transcription 1), and SNRPA (Small Nuclear Ribonucleoprotein Polypeptide A). These proteins have wide ranging roles in cellular processes such as cytoskeleton dynamics (PLEC), innate immune signaling (SYK and EIF2AK2), transcription factors (STAT1), and mRNA splicing and polyadenylation (SNRPA). 39/149 antiviral module proteins interacted with other antiviral module proteins, forming a total of 32 interactions. Many antiviral module proteins were indirectly linked in the network by interacting with a common first neighbor. For example, OAS1 physically associates with TRIM27, which physically associates with SNRPA (60–62). However, OAS1 and SNRPA do not directly physically associate with one another. Thus, changes to TRIM27 expression will may impact the antiviral response to ZIKV. Another example is that STAT1 physically associates with MAVS, and MAVS physically associates with OAS3 (63). Thus, although STAT1 and OAS3 do not physically associate with one another, they are connected through MAVS. The interconnectedness of the antiviral module network demonstrates how these antiviral molecules are part of a collective system that restricts ZIKV replication.

**Figure 7.**
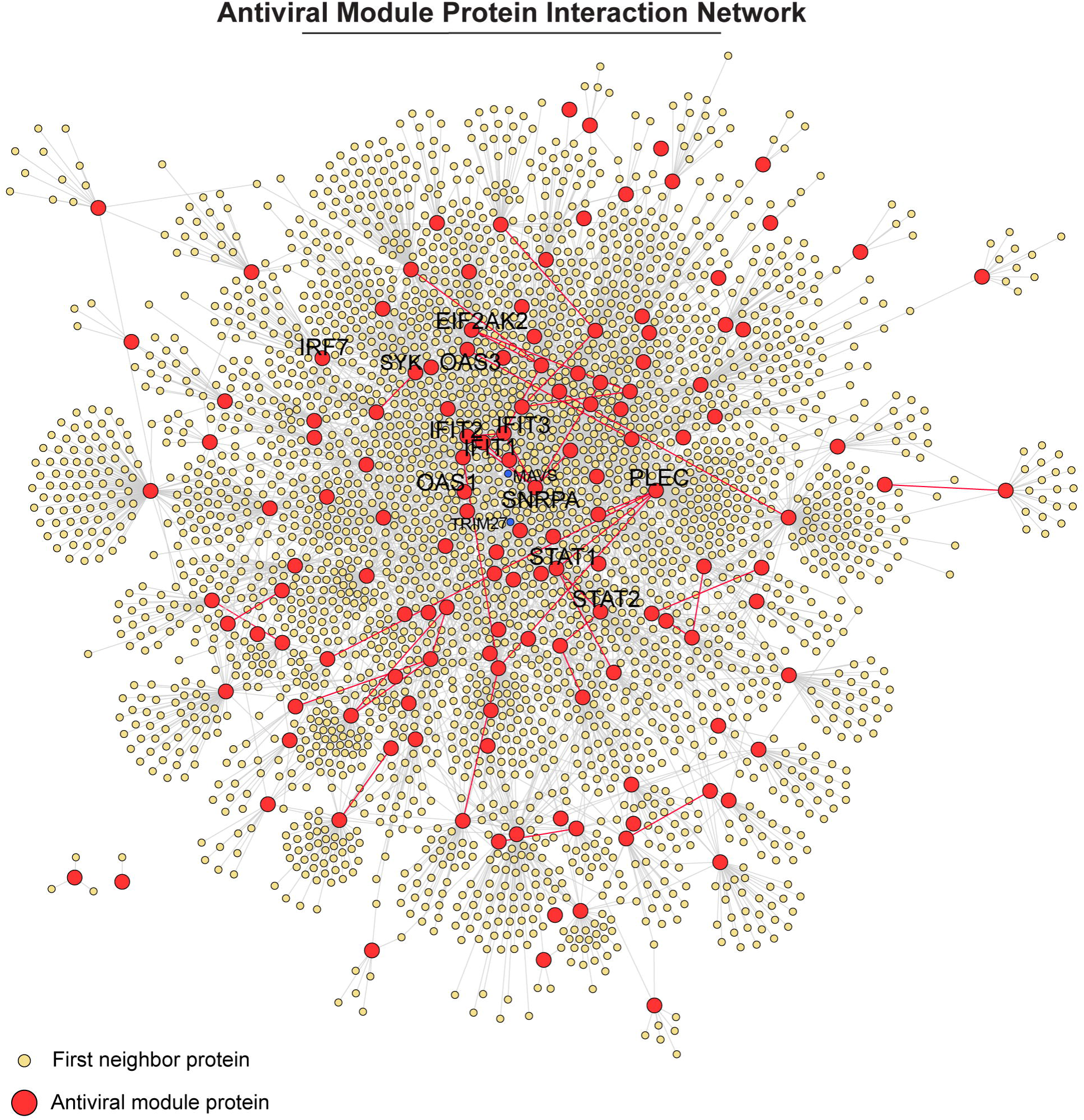
Protein-interaction network formed by antiviral module. Interaction network between human proteins encoded by the antiviral module genes and their first neighbors (proteins they interact with directly). The proteins are circular nodes (red = antiviral module protein, yellow = non-antiviral module protein) connected by edges (red = interactions between two antiviral module proteins, grey = all other interactions), which represent evidence-based protein interactions.

## Discussion

The host response to viral infection is often studied at the population level. This approach does not account for heterogeneity in the response to a virus across the spectrum of viral infection, from cells containing high viral loads to uninfected bystanders. In this work we aimed to investigate the transcriptional response to ZIKV within a population of human DCs with a low frequency of infection at the single cell level. To this end, we developed a ZIKV-inclusive scRNA-seq protocol that allowed us to distinguish between infected and bystander cells, characterize subpopulations within those and identify transcriptional modules that correlate with viral RNA. We present evidence that the antiviral response to ZIKV is a coordinated network of host genes that collectively restricts viral replication.

Previously, the host response to ZIKV infection has been investigated at the single cell level by flow cytometry (19, 64), mass cytometry (CyTOF) (40, 65), and microscopy (29, 66–68). Here we focused on the transcriptional response because scRNA-seq is inherently more sensitive than many other assays that rely upon antibody binding, it does not require a set of specific targets to be chosen prior to the experiment and is high-throughput and quantitative. Other flavivirus-inclusive assays have used the oligo-dT-primed SMART-seq2 assay, which involves index sorting cells into a plate to separate them (43, 49, 51–53). Here, we used a higher throughput droplet-based scRNA-seq assay which allows for profiling a higher number of cells and greater detection of rare cell types (69, 70). Including the ZIKV-specific primer increased the dynamic range of detecting viral RNA and allowed for more in-depth analysis of infected cells (*e.g.* exploration into infected subpopulations and correlation analyses between viral RNA and host gene expression). Using this approach, were able to more clearly mark cells that would otherwise reside at or below the limit of detection for infection. Our process for labeling infected and bystander cells is likely broadly applicable for all non-polyadenylated viruses.

During ZIKV infection in DCs, bystander cells took on an antiviral state rather than the infected cells, likely due to virus-mediated suppression of the response to IFN. The antiviral response in bystander cells has been observed in the context of a broad range of viruses such as respiratory syncytial virus (71), poxviruses (72), influenza virus (45), as well as in ZIKV infected macrophages (35). These results shed light on prior observations in human DCs, where ZIKV infection induced a robust expression of antiviral genes using bulk transcriptional analysis (19). The major transcriptional signatures observed in Bowen et al. were likely emanating from the bystander cells rather than ZIKV-infected cells. Population level analysis should be used with caution and an understanding that the ratio between infected and bystander cells may influence the measured output.

Type I IFN plays a critical role in the coordinated antiviral and inflammatory response during viral infection (73). In this study, *IFNB1* expression was exclusively found in infected cells, and within the population of infected cells *IFNB1* expression was rare. Considering the enriched type I IFN response we observed in bystander cells, the small number of IFN-β expressing cells had a clear effect. With less than 1% of the cells in the whole population expressing *IFNB1,* bystander cells expressed a substantial portion of the same differentially expressed genes in the group treated with soluble IFN-β. To our surprise, an even smaller fraction of DCs expressed *IFNA1* compared to *IFNB1*. IFN-α induction is canonically seen as a two-step process during viral infection, whereby *IFNB* expression leads to transcription and activation of *IRF7*, which is a master regulator of the IFN-α gene, leading to IFN-α gene expression (74, 75). In this study, IFN-β gene expression occurred only in the infected cells, while upregulated expression of *IRF7* occurred in the bystander cells in the context of ZIKV infection. The poor IFN-α induction in the bystander cells indicates that viral sensing by RLRs may be necessary for IRF-7 activation, similar to IRF-3 for induction of IFN-β. Conversely, *IRF7* expression negatively correlated with viral RNA, suggesting a ZIKV specific inhibition of *IRF7* expression. However, even in the infected cells that expressed *IRF7*, where RLR sensing should be present, the vast majority did not express *IFNA1*, suggesting that there may also be ZIKV mediated inhibition of IRF-7 activation or function. Another possibility is that there are other necessary co-factors for IFN-α induction such as the enhancesome for IFN-β induction (76). These results prompt the need for future studies to elucidate the disconnect between regulation of *IFNB1* and *IFNA1* in the context of ZIKV infection at the single cell level.

We observed an inverse relationship between viral RNA and a portion of host genes which we defined as the antiviral module by evaluating the transcription at the single cell level. In further support that this transcriptional module is antiviral, the uninfected bystander cells were also enriched for the module alongside infected cells with low viral RNA. Conversely, cells with high viral RNA had low antiviral module scores suggesting potential ZIKV mediated suppression. Indeed, prior studies have found that ZIKV nonstructural proteins antagonize antiviral pathways (Reviewed by Serman and Gack in 2019 (25)), including inhibition of TBK1 phosphorylation (27) and inhibition of JAK-STAT signaling via degradation JAK1 (28), inhibition of JAK1 and TYK2 activation (20), STAT2 degradation via the proteasome (26, 29, 32), and inhibition of STAT1 phosphorylation (19–21, 32). However, the vast majority of these studies were performed in the context of high multiplicity of infection or protein overexpression which may be less representative of physiologic conditions *in vivo* (77). We demonstrate here that ZIKV-host interactions can be evaluated in the context of a low frequency infection in primary cells using scRNA-seq. The use of our ZIKV-inclusive scRNA-seq protocol will be useful in further investigation into evasion of the antiviral response by specific ZIKV isolates.

In this work, we identified an antiviral network comprised of both IFN-dependent and independent genes that restrict ZIKV infection of DCs. There are hundreds of genes, commonly referred to as ISGs, that are induced upon activation of the type I IFN pathway, some of which have been found to be antiviral on an individual basis, yet insufficient to activate the full extent of the antiviral state alone (78–80). Given the need to protect against wide variety of pathogens and our observation that not every cell expresses the same set of ISGs or to the same magnitude under the same conditions, this redundancy may serve an evolutionary purpose (80). Prior work has shown that pairs of ISGs can work in an additive or synergistic antiviral manner (78, 81–83). Furthermore, there have been a few reports demonstrating that select ISGs with antiviral activity against an array of viruses form a protein-interaction network (63, 84, 85). Here, we expand on that concept and propose a new paradigm for the antiviral response to a specific virus, combining an unbiased list of genes that highly correlate with viral RNA on a per cell basis with experimental protein-interaction data. Future work should be done to evaluate this antiviral network in the context of ZIKV infection rather than individual genes.

One limitation of this study is that infected cells are defined independent of viral strand specificity. The primer used in the ZIKV-inclusive assay does not discriminate between positive strand, which is the genomic RNA strand, or negative, which is the replicative intermediate strand. Further development of this assay to specifically prime the negative strand of the flavivirus genome would improve confidence in infected cell labeling. Another limitation is that this study is based on transcriptional changes at single cell level. A single cell proteomic analysis such as outlined in Schoof et al. combined with transcriptomic analysis would provide a broader picture of the antiviral response during ZIKV infection (86).

In this study, we developed and implemented ZIKV-inclusive scRNA-seq with improved detection of ZIKV RNA and infected cells. Using this assay, we profiled the antiviral landscape in a population of human moDCs infected with ZIKV at the single cell level. We uncovered a network of genes that negatively correlate with viral RNA and that are enriched in bystander cells. Here we propose a new model of the antiviral response to ZIKV infection: a coordination of antiviral genes that collectively restrict ZIKV infection. These results lay the groundwork for further evaluation of the host response to flavivirus infection at the single cell level.

## Data availability

Single cell RNA sequencing data are publicly accessible through the Gene Expression Omnibus under accession number GSE230571. Code for single cell RNA sequencing analysis will be made available upon request.

## Acknowledgements

This work was supported by the Emory National Primate Research Center Flow Cytometry Core. Thank you to the Emory National Primate Research Center Genomics Core for performing the single cell RNA sequencing. Thank you to Jane Lawson for coordinating the healthy donors for acquisition of blood for PBMCs. This work was funded by the NIH R01 AI149486 and U01 AI131566.

## Supplemental Information Figure Legends

**Supplemental Figure 1.** T**i**trating **ZIKV-specific primer for 10x Genomics single cell RNA sequencing assay.** VeroE6 cells were infected with ZIKV. At 24 hours post-infection (hpi), mock and ZIKV-infected VeroE6 cells were mixed 1:1 and loaded onto the 10x Genomics Chromium Next GEM chip. The ZIKV-specific primer was introduced into the reverse transcription master mix for a final concentration of 0, 0.1, 1, and 10 µM. (A) Depth of sequencing for each group. (B) Schematic for the ZIKV-specific primer (left) and % of reads mapped to the ZIKV genome upstream of the ZIKV-specific primer annealing site (right). Violin plots displaying the expression levels of (C) ZIKV and (D) housekeeping genes across all groups.

**Supplemental Figure 2.** H**u**man **monocyte-derived dendritic cell flow cytometry.** Flow cytometry analysis of moDCs from cells used in scRNA-seq experiment at 24 hpi.

**Supplemental Figure 3.** H**e**atmap **of IFN-β signature genes.** Expression of top DEGs within the IFN-β treatment (termed IFN-β module) in mock, IFN-β, bystander, and ZIKV-infected cells. IFN-β treatment DEGs were filtered based on adj p-value < 0.05 and average log2 fold change > 1 in IFN-β treatment relative to mock. Data are represented as z-score. Colors: mock = green, IFN-β = blue, infected = red, bystander = yellow. DEGs = differentially expressed genes.

**Supplemental Figure 4.** T**o**tal **cells analyzed for transcriptional analysis.** Cells were subject to quality control filtering to remove low quality cells and doublets. These are the final numbers of cells included all downstream transcriptional analysis of VeroE6 and monocyte-derived dendritic cells (moDCs).

## Materials and Methods

### Ethics statement

Human peripheral blood monocular cells (PMBCs) were obtained from healthy donors in accordance with the Emory University Institutional review board according to IRB protocol IRB00045821.

### Cell lines

Cells were grown at 37°C with 5% CO2. VeroE6 cells (ATCC) and cultured in complete DMEM. Complete DMEM was prepared as follows: Dulbecco modified Eagle medium (DMEM; Lonza, Cat# 12-614Q supplemented with 10% heat-inactivated fetal bovine serum (FBS) and 1x Penicillin-Streptomycin (Corning, Cat# 30-002-CI).

### Generation of monocyte-derived dendritic cells

CD14+ monocytes were isolated from PMBCs of healthy human blood using the Mojosort^TM^ Human CD14 Selection Kit (BioLegend, Cat# 480026) To generate monocyte derived dendritic cells (moDCs), CD14+ monocytes were stimulated in non-tissue culture treated plates with 100ng/mL GM-CSF (R&D Systems, Cat# 7954-GM-020/CF) and IL-4 (R&D Systems, Cat# 6507-IL-025/CF) in complete RPMI. Complete RPMI was prepared as follows: RPMI 1640 (Corning, Cat# 10-040-CI) supplemented with 10% FBS, 2mM L-glutamine (Corning, Cat# 25-005-CI), 1mM HEPES (Corning, Cat# 25-060-CI), 1mM Sodium Pyruvate (Corning, Cat# 25-000-CI), 1x NEAA (Corning, Cat# 25-025-CI), and 1x Penicillin-Streptomycin. The next day, the media was replaced with fresh cytokines, removing un-adherent cells. After 4 more days in culture, differentiated moDCs consistently expressing CD14^−^ CD11c^+^ HLA-DR^+^ DC-SIGN^+^ were harvested in the supernatant. For experiments, moDCs were cultured in complete RPMI in U-bottom 96-well plates at 1e5 cells per well.

### Viruses

ZIKV strain PRVABC59 was used in all studies (34). VeroE6 cells were infected with ZIKV at MOI 5 and moDCs were infected with ZIKV at MOI 2.5 for 1 hr at 37°C. At 1 hpi, inoculum containing virus was removed and replaced with complete media, then cells were incubated for 24 hours. At 24 hpi, cells were treated with 0.25% trypsin-EDTA (Gibco, Cat# 25200-056) or Accutase® solution (Sigma-Aldrich, Cat# A6964) and the single cell suspension was harvested for flow cytometry analysis and scRNA-seq.

### Flow cytometry

Cells were blocked with Human TruStain FcX (BioLegend, Cat# 422302) in fluorescence activated cell sorting (FACS) buffer (1% FBS in 1x PBS) on ice for 10 minutes and then stained for viability (Ghost Dye^TM^ Red 780; Cell Signaling, Cat# 18452S) on ice for 20 minutes. moDCs were also stained for surface markers using BioLegend anti-human antibodies for monocyte and dendritic cell associated surface markers: CD14 (M5E2; BioLegend, Cat# 301814), CD11c (B-ly6; BioLegend, Cat# 563130), DC-SIGN (9E9A8; BioLegend, Cat# 330106), HLA-DR (G46-6; BioLegend, Cat# 562304), CD80 (2D10; BioLegend, Cat#305236), CD86 (IT2.2;

Biolegend, Cat# 305430). Cells were then washed with FACS buffer and fixed in 2% paraformaldehyde for 30 minutes. For intracellular ZIKV E protein staining, cells were incubated with 1x Transcription Factor Fix/Perm (Tonbo Biosciences, Cat# 1213-L150) for 20 minutes on ice and permeabilized by washing twice with 1x Flow Cytometry Perm Buffer (Tonbo Biosciences, Cat# 1213-L150). Pan-flavivirus envelope protein antibody 4G2 was conjugated to APC using Lightning-Link® APC Conjugation Kit (abcam, Cat# ab201807). Cells were then blocked with Human TruStain FcX in FACS on ice for 10 minutes followed by staining with 4G2-APC in 1x Flow Cytometry Perm Buffer on ice for 20 minutes. Cells were washed twice with 1x Flow Cytometry Perm Buffer and then resuspended in FACS buffer for flow cytometry. Samples were run on the LSR-II flow cytometry machine.

### Feature barcoding

Feature Barcoding was performed on moDCs prior to scRNA-seq to label mock and ZIKV treated cells. Cells were blocked with Human TruStain FcX on ice for 10 minutes and then surface stained with BioLegend antibodies (TotalSeq™-C0253 anti-human Hashtag 3 Antibody and TotalSeq™-C0252 anti-human Hashtag 2 Antibody) on ice for 20 minutes.

### Single cell RNA sequencing

Cells were washed with 0.04% w/v bovine serum albumin (BSA) in PBS and counted. The 10x Genomics Chromium Next GEM Single Cell 5’ v2 assay protocol (PN-1000264) was modified to be ZIKV-inclusive by adding a ZIKV-specific primer to the reverse transcription reaction mixture (**Figure 1**). All other steps were followed according to the manufacturer’s protocol. The ZIKV-specific primer 5’-<u>AAGCAGTGGTATCAACGCAGAGTAC</u>CCTTCCACAAAGTCCCTATTGC-3’) was previously published by Lanciotti et al. (55) and modified to contain the non-poly-dT tag (<u>underlined</u> <u>sequence</u>) for cDNA amplification. After bead cleanup, cDNA was amplified 13 cycles, and gene expression (5’ GEX) and feature barcoded libraries were generated using the 5’ v2 Library Construction Kit (PN-1000190). 5’ GEX libraries were constructed using cDNA that was first fragmented at 32°C for 5 min. Each sample was indexed with the Dual Index Kit TT Set A plate (PN-1000215) for 14 cycles of 98°C for 20s, 54°C for 30s, 72°C for 20s, with a final extension of 72°C for 1 min. Amplified feature barcoded libraries were constructed using DNA from supernatant cleanup. Each sample was indexed with the Dual Index Kit TN Set A plates (PN-1000250) for cycles of 98°C for 20s, 54°C for 30s, and 72°C for 20s, with a final extension of 72°C for 1 min.

For QC, final libraries un on a HS DNA chip on an Agilent Bioanalyzer 2100. The 5’GEX and feature barcoded libraries were pooled separately and then combined at an appropriate ratio in order to obtain the targeted read depth of 50,000/cell and 5,000/cell for the 5’ GEX and feature barcoded libraries, respectively. Samples were run on an Illumina NovaSeq S4 single lane (2-2.5 billion reads/run) at the Emory National Primate Research Center Genomics Core.

### Bioinformatics pipeline

A composite genome reference was created by concatenating the sequence files for both the host genomes and the ZIKV PRABC59 genome (KU501212.1). Features were added to the composite annotation noting the location of the primer sequence and regions up and downstream of that location. These composite reference files were used to create a 10x Cell Ranger reference that was used for alignment and processing. Data analysis was performed using the R package Seurat (v4.1). The Feature Barcode sequences in the moDC data were added as a hashtag oligo (HTO) assay and then normalized by a centered log ratio transformation. Feature Barcode sequences were demultiplexed using the HTODemux function in Seurat, selecting for the 0.9 positive quantile and singlets (i.e. cells with only either mock or ZIKV associated barcodes). Then moDCs were filtered to remove cells with >60,000 and <500 RNA counts, >1,000 unique genes (features), >25% mitochondrial genes, and >15% ribosomal genes. VeroE6 data were filtered to remove cells with >15,000 and <100 RNA counts, >5,000 unique genes (features), >2% mitochondrial genes, and >15% and <2% ribosomal genes. Filtered read counts were normalized scaled by a factor of 10,000 and natural log transformed (default settings for NormalizeData in Seurat). Doublets were computationally predicted using the R package DoubletFinder (v2.0.3) and subsequently removed. The optimal number of components (33) was inferred using the Elbow method and was used for UMAP dimension reduction. A total 15,808 moDCs and 39,035 VeroE6 cells were successfully captured and profiled for transcriptomic analysis (**Supplemental Figure 4**). Viral RNA was quantitated based on reads that mapped upstream of the ZIKV-specific primer between 0-996 nucleotides in the genome.

### Statistics

Gene expression is represented as natural log normalized values in all UMAP and violin plots unless otherwise stated. Pairwise comparisons between groups for expression of specific genes (**Figure 2**) were performed using the FindMarkers function in Seurat using the non-parametric Wilcoxon Rank Sum test and the Bonferroni-adjusted p-values were reported. After **Figure 2**, only samples with the ZIKV-specific primer added were analyzed. Gene Set Enrichment Analysis (GSEA) was performed with the R package fgsea (v1.20.0) with ranking based on average log2 fold change and 10,000 permutations. Gene expression data for all genes relative to mock used for GSEA was generated using FindMarkers function in Seurat with the following thresholds: min.pct = 0.01 and log2 fold change = 0.01. Hallmark gene sets from the Human Molecular Signatures Database (MSigDB) were used for GSEA (87, 88). UMAPs featuring gene set Hallmark pathways was performed using the sample level enrichment analysis (SLEA) method. Differential gene expression analysis (**Figure 5-6**) was performed with the FindMarkers function in Seurat using the non-parametric Wilcoxon Rank Sum test on genes expressed in a minimum fraction of 0.1 cells (min.pct = 0.1; default). Top differentially expressed genes (DEGs) were additionally filtered for Bonferroni-adjusted p-values <0.05.

### Protein-interaction network analysis

Antiviral module genes were converted to their protein counterparts and known protein-protein interactions were identified between antiviral proteins and their first neighbors through IntAct Molecular Interaction Database Application in Cytoscape (version 3.8.2)(89, 90). Results from IntAct were filtered for only human proteins. The Allegro Spring-Electric Algorithm was used to generate the network map layout (Allegro Layout Application, version 2.2.2). Duplicate edges and self-loops were removed from the network map in **Figure 7**. See Supplemental information for a table listing the IntAct Accenssion code, interaction type, detection method, and PubMed ID associated with each interaction pair.

## References

1. Dick GW, Kitchen SF, Haddow AJ. Zika virus. I. Isolations and serological specificity. Trans R Soc Trop Med Hyg. 1952;46(5):509–20.

2. Weaver SC, Costa F, Garcia-Blanco MA, Ko AI, Ribeiro GS, Saade G, et al. Zika virus: History, emergence, biology, and prospects for control. Antiviral research. 2016;130:69–80.

3. Brasil PcaPJPaMMEaRNRMaDLaWMaRRSaV. Zika Virus Infection in Pregnant Women in Rio de Janeiro. New England Journal of Medicine. 2016;375(24):2321–34.

4. Suthar MS, Ma DY, Thomas S, Lund JM, Zhang N, Daffis S, et al. IPS-1 is essential for the control of West Nile virus infection and immunity. PLoS Pathog. 2010;6(2):e1000757.

5. Ma DY, Suthar MS, Kasahara S, Gale M, Jr., Clark EA. CD22 is required for protection against West Nile virus Infection. J Virol. 2013;87(6):3361–75.

6. Pinto AK, Ramos HJ, Wu X, Aggarwal S, Shrestha B, Gorman M, et al. Deficient IFN signaling by myeloid cells leads to MAVS-dependent virus-induced sepsis. PLoS Pathog. 2014;10(4):e1004086.

7. Aye KS, Charngkaew K, Win N, Wai KZ, Moe K, Punyadee N, et al. Pathologic highlights of dengue hemorrhagic fever in 13 autopsy cases from Myanmar. Hum Pathol. 2014;45(6):1221–33.

8. Balsitis SJ, Coloma J, Castro G, Alava A, Flores D, McKerrow JH, et al. Tropism of dengue virus in mice and humans defined by viral nonstructural protein 3-specific immunostaining. Am J Trop Med Hyg. 2009;80(3):416–24.

9. Wu SJ, Grouard-Vogel G, Sun W, Mascola JR, Brachtel E, Putvatana R, et al. Human skin Langerhans cells are targets of dengue virus infection. Nat Med. 2000;6(7):816–20.

10. Barba-Spaeth G, Longman RS, Albert ML, Rice CM. Live attenuated yellow fever 17D infects human DCs and allows for presentation of endogenous and recombinant T cell epitopes. J Exp Med. 2005;202(9):1179–84.

11. Brandler S, Brown N, Ermak TH, Mitchell F, Parsons M, Zhang Z, et al. Replication of chimeric yellow fever virus-dengue serotype 1-4 virus vaccine strains in dendritic and hepatic cells. Am J Trop Med Hyg. 2005;72(1):74–81.

12. Michlmayr D, Andrade P, Gonzalez K, Balmaseda A, Harris E. CD14+CD16+ monocytes are the main target of Zika virus infection in peripheral blood mononuclear cells in a paediatric study in Nicaragua. Nat Microbiol. 2017;2(11):1462–70.

13. Silveira ELV, Rogers KA, Gumber S, Amancha P, Xiao P, Woollard SM, et al. Immune Cell Dynamics in Rhesus Macaques Infected with a Brazilian Strain of Zika Virus. J Immunol. 2017;199(3):1003–11.

14. O’Connor MA, Tisoncik-Go J, Lewis TB, Miller CJ, Bratt D, Moats CR, et al. Early cellular innate immune responses drive Zika viral persistence and tissue tropism in pigtail macaques. Nat Commun. 2018;9(1):3371.

15. Hamel R, Dejarnac O, Wichit S, Ekchariyawat P, Neyret A, Luplertlop N, et al. Biology of Zika Virus Infection in Human Skin Cells. J Virol. 2015;89(17):8880–96.

16. Reynoso GV, Gordon DN, Kalia A, Aguilar CC, Malo CS, Aleshnick M, et al. Zika virus spreads through infection of lymph node-resident macrophages. Cell Rep. 2023;42(2):112126.

17. Österlund P, Jiang M, Westenius V, Kuivanen S, Järvi R, Kakkola L, et al. Asian and African lineage Zika viruses show differential replication and innate immune responses in human dendritic cells and macrophages. Sci Rep. 2019;9(1):15710.

18. Vielle NJ, Zumkehr B, García-Nicolás O, Blank F, Stojanov M, Musso D, et al. Silent infection of human dendritic cells by African and Asian strains of Zika virus. Sci Rep. 2018;8(1):5440.

19. Bowen JR, Quicke KM, Maddur MS, O’Neal JT, McDonald CE, Fedorova NB, et al. Zika Virus Antagonizes Type I Interferon Responses during Infection of Human Dendritic Cells. PLoS Pathog. 2017;13(2):e1006164.

20. Zimmerman MG, Bowen JR, McDonald CE, Young E, Baric RS, Pulendran B, et al. STAT5: a Target of Antagonism by Neurotropic Flaviviruses. J Virol. 2019;93(23).

21. Yang D, Chu H, Lu G, Shuai H, Wang Y, Hou Y, et al. STAT2-dependent restriction of Zika virus by human macrophages but not dendritic cells. Emerg Microbes Infect. 2021;10(1):1024–37.

22. Branche E, Wang YT, Viramontes KM, Valls Cuevas JM, Xie J, Ana-Sosa-Batiz F, et al. SREBP2-dependent lipid gene transcription enhances the infection of human dendritic cells by Zika virus. Nat Commun. 2022;13(1):5341.

23. Park T, Kang MG, Baek SH, Lee CH, Park D. Zika virus infection differentially affects genome-wide transcription in neuronal cells and myeloid dendritic cells. PLoS One. 2020;15(4):e0231049.

24. Sun X, Hua S, Chen HR, Ouyang Z, Einkauf K, Tse S, et al. Transcriptional Changes during Naturally Acquired Zika Virus Infection Render Dendritic Cells Highly Conducive to Viral Replication. Cell Rep. 2017;21(12):3471–82.

25. Serman TM, Gack MU. Evasion of Innate and Intrinsic Antiviral Pathways by the Zika Virus. Viruses. 2019;11(10).

26. Grant A, Ponia SS, Tripathi S, Balasubramaniam V, Miorin L, Sourisseau M, et al. Zika Virus Targets Human STAT2 to Inhibit Type I Interferon Signaling. Cell Host Microbe. 2016;19(6):882–90.

27. Xia H, Luo H, Shan C, Muruato AE, Nunes BTD, Medeiros DBA, et al. An evolutionary NS1 mutation enhances Zika virus evasion of host interferon induction. Nat Commun. 2018;9(1):414.

28. Wu Y, Liu Q, Zhou J, Xie W, Chen C, Wang Z, et al. Zika virus evades interferon-mediated antiviral response through the co-operation of multiple nonstructural proteins in vitro. Cell Discov. 2017;3:17006.

29. Kumar A, Hou S, Airo AM, Limonta D, Mancinelli V, Branton W, et al. Zika virus inhibits type-I interferon production and downstream signaling. EMBO Rep. 2016;17(12):1766–75.

30. Li W, Li N, Dai S, Hou G, Guo K, Chen X, et al. Zika virus circumvents host innate immunity by targeting the adaptor proteins MAVS and MITA. Faseb j. 2019;33(9):9929–44.

31. Fanunza E, Grandi N, Quartu M, Carletti F, Ermellino L, Milia J, et al. INMI1 Zika Virus NS4B Antagonizes the Interferon Signaling by Suppressing STAT1 Phosphorylation. Viruses. 2021;13(12).

32. Hertzog J, Dias Junior AG, Rigby RE, Donald CL, Mayer A, Sezgin E, et al. Infection with a Brazilian isolate of Zika virus generates RIG-I stimulatory RNA and the viral NS5 protein blocks type I IFN induction and signaling. Eur J Immunol. 2018;48(7):1120–36.

33. Zimmerman MG, Quicke KM, O’Neal JT, Arora N, Machiah D, Priyamvada L, et al. Cross-Reactive Dengue Virus Antibodies Augment Zika Virus Infection of Human Placental Macrophages. Cell Host Microbe. 2018;24(5):731–42.e6.

34. Quicke KM, Bowen JR, Johnson EL, McDonald CE, Ma H, O’Neal JT, et al. Zika Virus Infects Human Placental Macrophages. Cell Host Microbe. 2016;20(1):83–90.

35. Carlin AF, Vizcarra EA, Branche E, Viramontes KM, Suarez-Amaran L, Ley K, et al. Deconvolution of pro- and antiviral genomic responses in Zika virus-infected and bystander macrophages. Proc Natl Acad Sci U S A. 2018;115(39):E9172–e81.

36. Hamlin RE, Rahman A, Pak TR, Maringer K, Mena I, Bernal-Rubio D, et al. High-dimensional CyTOF analysis of dengue virus-infected human DCs reveals distinct viral signatures. JCI Insight. 2017;2(13).

37. Kelly JN, Laloli L, V’Kovski P, Holwerda M, Portmann J, Thiel V, et al. Comprehensive single cell analysis of pandemic influenza A virus infection in the human airways uncovers cell-type specific host transcriptional signatures relevant for disease progression and pathogenesis. Front Immunol. 2022;13:978824.

38. Sun J, Vera JC, Drnevich J, Lin YT, Ke R, Brooke CB. Single cell heterogeneity in influenza A virus gene expression shapes the innate antiviral response to infection. PLoS Pathog. 2020;16(7):e1008671.

39. Hancock AS, Stairiker CJ, Boesteanu AC, Monzón-Casanova E, Lukasiak S, Mueller YM, et al. Transcriptome Analysis of Infected and Bystander Type 2 Alveolar Epithelial Cells during Influenza A Virus Infection Reveals In Vivo Wnt Pathway Downregulation. J Virol. 2018;92(21).

40. Fenutria R, Maringer K, Potla U, Bernal-Rubio D, Evans MJ, Harris E, et al. CyTOF Profiling of Zika and Dengue Virus-Infected Human Peripheral Blood Mononuclear Cells Identifies Phenotypic Signatures of Monotype Subsets and Upregulation of the Interferon-Inducible Protein CD169. mSphere. 2021;6(3):e0050521.

41. Cristinelli S, Ciuffi A. The use of single-cell RNA-Seq to understand virus-host interactions. Curr Opin Virol. 2018;29:39–50.

42. Zanini FaPS-YaBEaESaQSR. Single-cell transcriptional dynamics of flavivirus infection. eLife. 2018;7:e32942, citation = eLife 2018;7 e.

43. O’Neal JT, Upadhyay AA, Wolabaugh A, Patel NB, Bosinger SE, Suthar MS. West Nile Virus-Inclusive Single-Cell RNA Sequencing Reveals Heterogeneity in the Type I Interferon Response within Single Cells. J Virol. 2019;93(6).

44. Russell AB, Trapnell C, Bloom JD. Extreme heterogeneity of influenza virus infection in single cells. Elife. 2018;7.

45. Steuerman Y, Cohen M, Peshes-Yaloz N, Valadarsky L, Cohn O, David E, et al. Dissection of Influenza Infection In Vivo by Single-Cell RNA Sequencing. Cell Syst. 2018;6(6):679–91.e4.

46. Onorati M, Li Z, Liu F, Sousa André MM, Nakagawa N, Li M, et al. Zika Virus Disrupts Phospho-TBK1 Localization and Mitosis in Human Neuroepithelial Stem Cells and Radial Glia. Cell Reports. 2016;16(10):2576–92.

47. Gorman MJ, Caine EA, Zaitsev K, Begley MC, Weger-Lucarelli J, Uccellini MB, et al. An Immunocompetent Mouse Model of Zika Virus Infection. Cell Host Microbe. 2018;23(5):672–85.e6.

48. Wells MF, Nemesh J, Ghosh S, Mitchell JM, Salick MR, Mello CJ, et al. Natural variation in gene expression and viral susceptibility revealed by neural progenitor cell villages. Cell Stem Cell. 2023;30(3):312–32.e13.

49. Zanini F, Pu S-Y, Bekerman E, Einav S, Quake SR. Single-cell transcriptional dynamics of flavivirus infection. eLife. 2018;7:7:e32942.

50. Yang W, Liu L-B, Liu F-L, Wu Y-H, Zhen Z-D, Fan D-Y, et al. Single-cell RNA sequencing reveals the fragility of male spermatogenic cells to Zika virus-induced complement activation. Nature Communications. 2023;14(1):2476.

51. Picelli S. Single-cell RNA-sequencing: The future of genome biology is now. RNA Biol. 2017;14(5):637–50.

52. Sanborn MA, Li T, Victor K, Siegfried H, Fung C, Rothman AL, et al. Analysis of cell-associated DENV RNA by oligo(dT) primed 5’ capture scRNAseq. Sci Rep. 2020;10(1):9047.

53. Zanini F, Robinson ML, Croote D, Sahoo MK, Sanz AM, Ortiz-Lasso E, et al. Virus- inclusive single-cell RNA sequencing reveals the molecular signature of progression to severe dengue. Proc Natl Acad Sci U S A. 2018;115(52):E12363–e9.

54. . !!! INVALID CITATION !!! (41, 45).

55. Lanciotti RS, Kosoy OL, Laven JJ, Velez JO, Lambert AJ, Johnson AJ, et al. Genetic and serologic properties of Zika virus associated with an epidemic, Yap State, Micronesia, 2007. Emerg Infect Dis. 2008;14(8):1232–9.

56. Ratnasiri K, Wilk AJ, Lee MJ, Khatri P, Blish CA. Single-cell RNA-seq methods to interrogate virus-host interactions. Seminars in Immunopathology. 2023;45(1):71–89.

57. Zhao M, Zhang J, Phatnani H, Scheu S, Maniatis T. Stochastic expression of the interferon-β gene. PLoS Biol. 2012;10(1):e1001249.

58. Hare DN, Baid K, Dvorkin-Gheva A, Mossman KL. Virus-Intrinsic Differences and Heterogeneous IRF3 Activation Influence IFN-Independent Antiviral Protection. iScience. 2020;23(12):101864.

59. Zawatzky R, De Maeyer E, De Maeyer-Guignard J. Identification of individual interferon-producing cells by in situ hybridization. Proc Natl Acad Sci U S A. 1985;82(4):1136–40.

60. Fragoza R, Das J, Wierbowski SD, Liang J, Tran TN, Liang S, et al. Extensive disruption of protein interactions by genetic variants across the allele frequency spectrum in human populations. Nat Commun. 2019;10(1):4141.

61. Rolland T, Taşan M, Charloteaux B, Pevzner SJ, Zhong Q, Sahni N, et al. A proteome-scale map of the human interactome network. Cell. 2014;159(5):1212–26.

62. Luck K, Kim DK, Lambourne L, Spirohn K, Begg BE, Bian W, et al. A reference map of the human binary protein interactome. Nature. 2020;580(7803):402–8.

63. Li S, Wang L, Berman M, Kong YY, Dorf ME. Mapping a dynamic innate immunity protein interaction network regulating type I interferon production. Immunity. 2011;35(3):426–40.

64. Michlmayr D, Andrade P, Gonzalez K, Balmaseda A, Harris E. CD14 + CD16 + monocytes are the main target of Zika virus infection in peripheral blood mononuclear cells in a paediatric study in Nicaragua. Nat Microbiol. 2017;2(11):1462–70.

65. Michlmayr D, Kim EY, Rahman AH, Raghunathan R, Kim-Schulze S, Che Y, et al. Comprehensive Immunoprofiling of Pediatric Zika Reveals Key Role for Monocytes in the Acute Phase and No Effect of Prior Dengue Virus Infection. Cell Rep. 2020;31(4):107569.

66. Monel B, Compton AA, Bruel T, Amraoui S, Burlaud-Gaillard J, Roy N, et al. Zika virus induces massive cytoplasmic vacuolization and paraptosis-like death in infected cells. Embo j. 2017;36(12):1653–68.

67. Long RKM, Moriarty KP, Cardoen B, Gao G, Vogl AW, Jean F, et al. Super resolution microscopy and deep learning identify Zika virus reorganization of the endoplasmic reticulum. Sci Rep. 2020;10(1):20937.

68. Cortese M, Goellner S, Acosta EG, Neufeldt CJ, Oleksiuk O, Lampe M, et al. Ultrastructural Characterization of Zika Virus Replication Factories. Cell Rep. 2017;18(9):2113–23.

69. Baran-Gale J, Chandra T, Kirschner K. Experimental design for single-cell RNA sequencing. Brief Funct Genomics. 2018;17(4):233–9.

70. Wang X, He Y, Zhang Q, Ren X, Zhang Z. Direct Comparative Analyses of 10X Genomics Chromium and Smart-seq2. Genomics Proteomics Bioinformatics. 2021;19(2):253–66.

71. Czerkies M, Kochańczyk M, Korwek Z, Prus W, Lipniacki T. Respiratory Syncytial Virus Protects Bystander Cells against Influenza A Virus Infection by Triggering Secretion of Type I and Type III Interferons. J Virol. 2022;96(22):e0134122.

72. Melo-Silva CR, Roman MI, Knudson CJ, Tang L, Xu RH, Tassetto M, et al. Interferon partly dictates a divergent transcriptional response in poxvirus-infected and bystander inflammatory monocytes. Cell Rep. 2022;41(8):111676.

73. Shalek AK, Satija R, Shuga J, Trombetta JJ, Gennert D, Lu D, et al. Single-cell RNA-seq reveals dynamic paracrine control of cellular variation. Nature. 2014;510(7505):363-9.

74. Honda K, Yanai H, Negishi H, Asagiri M, Sato M, Mizutani T, et al. IRF-7 is the master regulator of type-I interferon-dependent immune responses. Nature. 2005;434(7034):772-7.

75. Lazear HM, Lancaster A, Wilkins C, Suthar MS, Huang A, Vick SC, et al. IRF-3, IRF-5, and IRF-7 coordinately regulate the type I IFN response in myeloid dendritic cells downstream of MAVS signaling. PLoS Pathog. 2013;9(1):e1003118.

76. Apostolou E, Thanos D. Virus Infection Induces NF-kappaB-dependent interchromosomal associations mediating monoallelic IFN-beta gene expression. Cell. 2008;134(1):85–96.

77. Meischel T, Fritzlar S, Villalón-Letelier F, Smith JM, Brooks AG, Reading PC, et al. Caveats of Using Overexpression Approaches to Screen Cellular Host IFITM Proteins for Antiviral Activity. Pathogens. 2023;12(4).

78. Schoggins JW, Wilson SJ, Panis M, Murphy MY, Jones CT, Bieniasz P, et al. A diverse range of gene products are effectors of the type I interferon antiviral response. Nature. 2011;472(7344):481-5.

79. Zhou A, Paranjape JM, Der SD, Williams BR, Silverman RH. Interferon action in triply deficient mice reveals the existence of alternative antiviral pathways. Virology. 1999;258(2):435–40.

80. Schneider WM, Chevillotte MD, Rice CM. Interferon-stimulated genes: a complex web of host defenses. Annu Rev Immunol. 2014;32:513–45.

81. Karki S, Li MM, Schoggins JW, Tian S, Rice CM, MacDonald MR. Multiple interferon stimulated genes synergize with the zinc finger antiviral protein to mediate anti-alphavirus activity. PLoS One. 2012;7(5):e37398.

82. MacDonald MR, Machlin ES, Albin OR, Levy DE. The zinc finger antiviral protein acts synergistically with an interferon-induced factor for maximal activity against alphaviruses. J Virol. 2007;81(24):13509–18.

83. Li MM, Lau Z, Cheung P, Aguilar EG, Schneider WM, Bozzacco L, et al. TRIM25 Enhances the Antiviral Action of Zinc-Finger Antiviral Protein (ZAP). PLoS Pathog. 2017;13(1):e1006145.

84. Hubel P, Urban C, Bergant V, Schneider WM, Knauer B, Stukalov A, et al. A protein-interaction network of interferon-stimulated genes extends the innate immune system landscape. Nat Immunol. 2019;20(4):493–502.

85. Navratil V, de Chassey B, Meyniel L, Pradezynski F, André P, Rabourdin-Combe C, et al. System-level comparison of protein-protein interactions between viruses and the human type I interferon system network. J Proteome Res. 2010;9(7):3527–36.

86. Schoof EM, Furtwängler B, Üresin N, Rapin N, Savickas S, Gentil C, et al. Quantitative single-cell proteomics as a tool to characterize cellular hierarchies. Nat Commun. 2021;12(1):3341.

87. Subramanian A, Tamayo P, Mootha VK, Mukherjee S, Ebert BL, Gillette MA, et al. Gene set enrichment analysis: a knowledge-based approach for interpreting genome-wide expression profiles. Proc Natl Acad Sci U S A. 2005;102(43):15545–50.

88. Mootha VK, Lindgren CM, Eriksson KF, Subramanian A, Sihag S, Lehar J, et al. PGC-1alpha-responsive genes involved in oxidative phosphorylation are coordinately downregulated in human diabetes. Nat Genet. 2003;34(3):267–73.

89. Orchard S, Ammari M, Aranda B, Breuza L, Briganti L, Broackes-Carter F, et al. The MIntAct project--IntAct as a common curation platform for 11 molecular interaction databases. Nucleic Acids Res. 2014;42(Database issue):D358–63.

90. Shannon P, Markiel A, Ozier O, Baliga NS, Wang JT, Ramage D, et al. Cytoscape: a software environment for integrated models of biomolecular interaction networks. Genome Res. 2003;13(11):2498–504.

