## Supplemental Figures for "Single cell analysis reveals an antiviral network that controls Zika virus infection in human dendritic cells"

### Figure S1

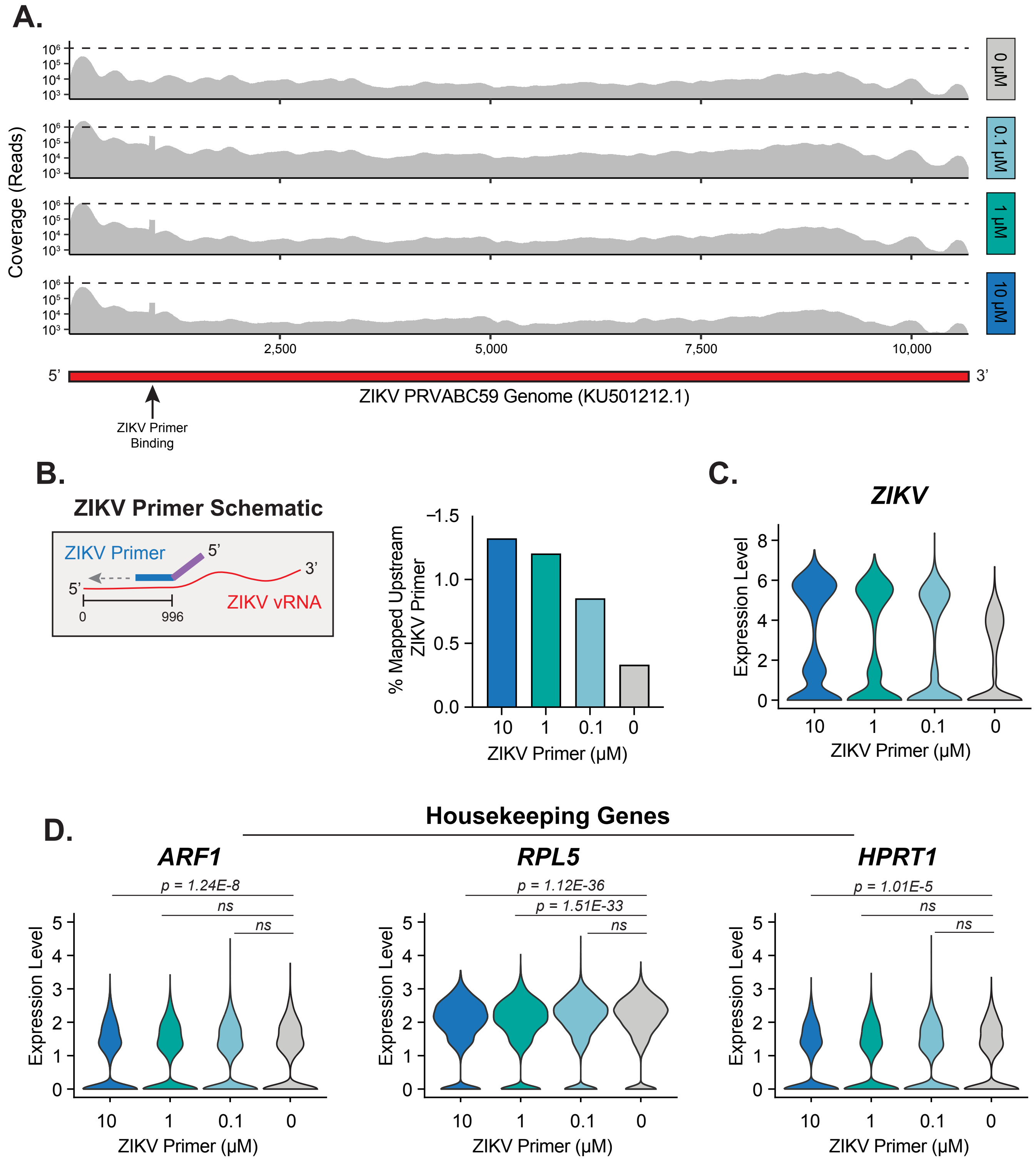
A.
Coverage (Reads)
10⁶
10⁵
10⁴
10³
0 µM
0.1 µM
1 µM
10 µM
2,500
5,000
7,500
10,000
5'
3'
ZIKV PRVABC59 Genome (KU501212.1)
ZIKV Primer Binding
B.
ZIKV Primer Schematic
ZIKV Primer
5'
3'
5'
ZIKV vRNA
0
996
% Mapped Upstream ZIKV Primer
−1.5
1.0
0.5
0.0
10
1
0.1
0
ZIKV Primer (µM)
C.
ZIKV
Expression Level
8
6
4
2
0
10
1
0.1
0
ZIKV Primer (µM)
D.
Housekeeping Genes
ARF1
p = 1.24E-8
ns
ns
Expression Level
ZIKV Primer (µM)
RPL5
p = 1.12E-36
p = 1.51E-33
ns
Expression Level
ZIKV Primer (µM)
HPRT1
p = 1.01E-5
ns
ns
Expression Level
ZIKV Primer (µM)

### Figure S2

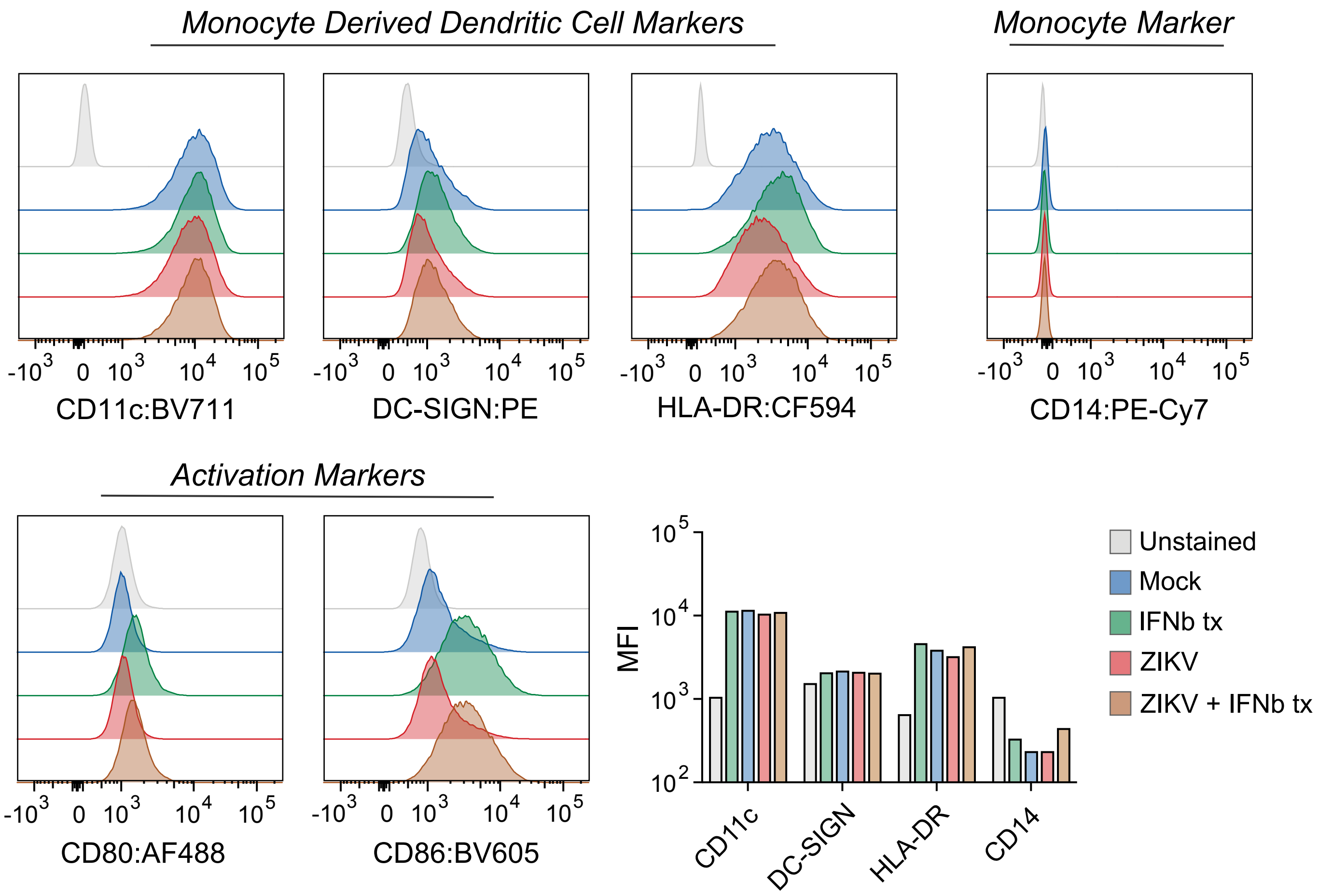
Monocyte Derived Dendritic Cell Markers
Monocyte Marker
CD11c:BV711
DC-SIGN:PE
HLA-DR:CF594
CD14:PE-Cy7
Activation Markers
CD80:AF488
CD86:BV605
MFI
CD11c
DC-SIGN
HLA-DR
CD14
Unstained
Mock
IFNb tx
ZIKV
ZIKV + IFNb tx

### Figure S3

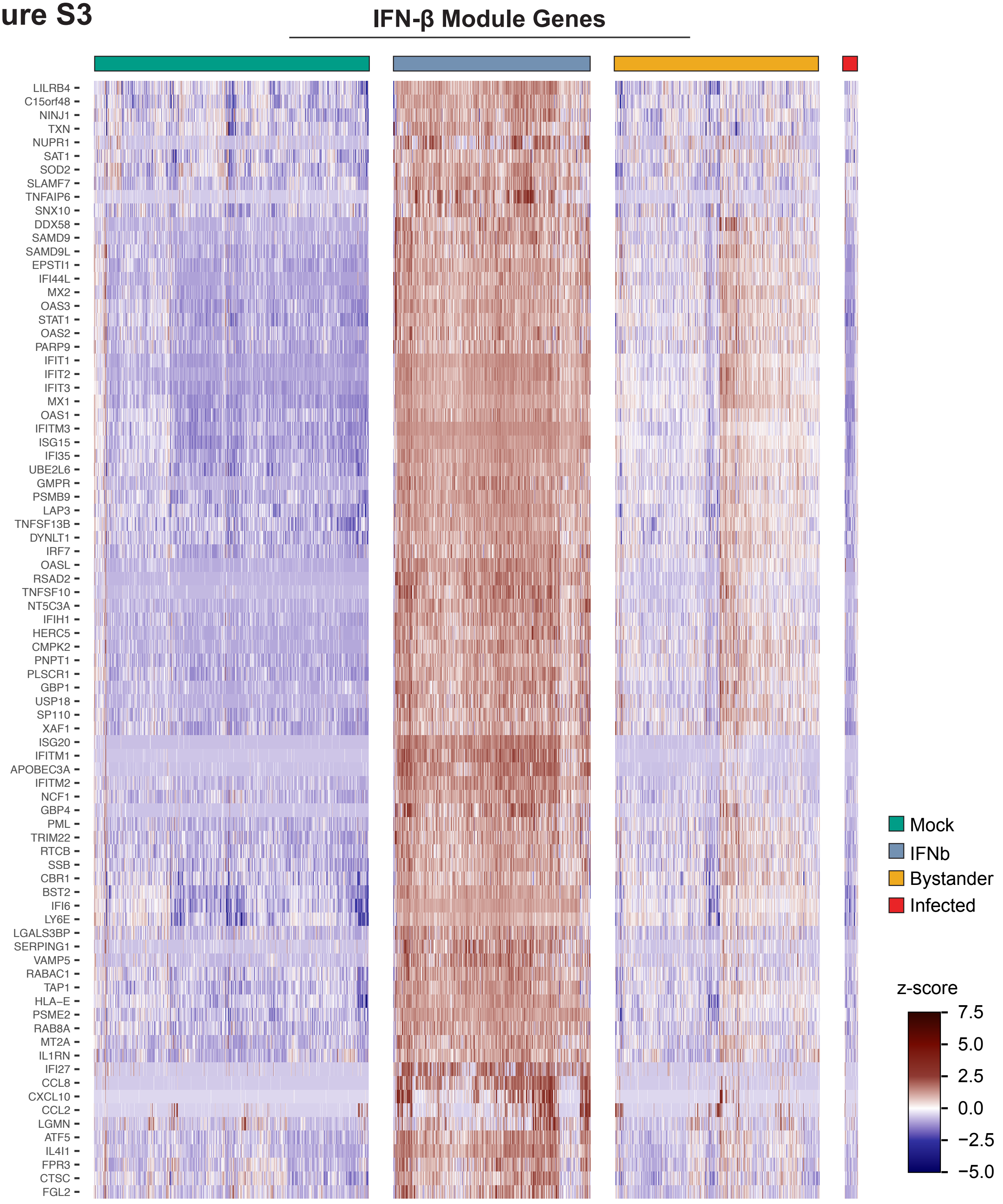
IFN-β Module Genes
LILRB4
C15orf48
NINJ1
TXN
NUPR1
SAT1
SOD2
SLAMF7
TNFAIP6
SNX10
DDX58
SAMD9
SAMD9L
EPSTI1
IFI44L
MX2
OAS3
STAT1
OAS2
PARP9
IFIT1
IFIT2
IFIT3
MX1
OAS1
IFITM3
ISG15
IFI35
UBE2L6
GMPR
PSMB9
LAP3
TNFSF13B
DYNLT1
IRF7
OASL
RSAD2
TNFSF10
NT5C3A
IFIH1
HERC5
CMPK2
PNPT1
PLSCR1
GBP1
USP18
SP110
XAF1
ISG20
IFITM1
APOBEC3A
IFITM2
NCF1
GBP4
PML
TRIM22
RTCB
SSB
CBR1
BST2
IFI6
LY6E
LGALS3BP
SERPING1
VAMP5
RABAC1
TAP1
HLA−E
PSME2
RAB8A
MT2A
IL1RN
IFI27
CCL8
CXCL10
CCL2
LGMN
ATF5
IL4I1
FPR3
CTSC
FGL2
Mock
IFNb
Bystander
Infected
z-score
7.5
5.0
2.5
0.0
−2.5
−5.0

### Figure S5

#### Monocyte-derived dendritic cells

| SampleID | Group | n_cells |
|---|---|---|
| 1 | IFNb tx | 2042 |
| 1 | Mock | 1733 |
| 2 | Bystander | 2117 |
| 2 | Infected | 136 |
| 2 | Mock | 1108 |
| 2 | Dump* | 482 |
| 3 | Bystander/IFNb tx | 2635 |
| 3 | Infected | 1 |
| 3 | Mock | 1258 |
| 3 | Dump* | 44 |
| 4 | Bystander | 2737 |
| 4 | Infected | 146 |
| 4 | Mock | 1283 |
| 4 | Dump* | 86 |
| | Total | 15808 |

Dump* = ZIKV: ZIKV 0-1; Mock: ZIKV > 0; cells removed from analysis

#### VeroE6 cells

| SampleID | Group | n_cells |
|---|---|---|
| 1 | ZIKV Primer, 10 uM | 10197 |
| 2 | ZIKV Primer, 1 uM | 9673 |
| 3 | ZIKV Primer, 0.1 uM | 9348 |
| 4 | ZIKV Primer, 0 uM | 9817 |
| | Total | 39035 |
